# Inhibition of Cxcr4 chemokine receptor signaling improves habituation learning and increases cAMP-PKA signaling in a zebrafish model of Neurofibromatosis type 1

**DOI:** 10.1101/2025.05.30.657071

**Authors:** Andrew H. Miller, Yeng Yang, Natalie Schmidt, Jaffna Mathiaparanam, Mark E. Berres, Mary C. Halloran

## Abstract

Neurofibromatosis type 1 (NF1) is a neurogenetic disorder caused by loss of function mutations in the gene neurofibromin 1 (*NF1*). *NF1* encodes neurofibromin, a multifunctional tumor suppressing protein that regulates Ras, cAMP, and dopamine signaling. NF1 predisposes patients to a wide range of symptoms, including peripheral nerve tumors, brain tumors, and cognitive dysfunction. Despite considerable work using animal models to investigate the role of neurofibromin in behavior, translating research into treatment for NF1-associated cognitive dysfunction has not yet been successful. Here, we identify Cxcr4 chemokine receptor signaling as a regulator of habituation learning and modulator of cAMP-PKA signaling in *nf1* mutant larval zebrafish. Combining a small-molecule drug screen and RNAseq analysis, we show that *cxcr4b* expression is increased in *nf1* mutants and that pharmacological inhibition of Cxcr4 with AMD3100 (Plerixafor) improves habituation learning in *nf1* mutants. We further demonstrate that Plerixafor treatment activates cAMP-PKA pathway signaling but has limited effects on Ras-Raf-MEK-ERK pathway signaling in the *nf1* mutant brain. CXCR4 was previously identified as a potential therapeutic target for neurofibromin-deficient tumorigenesis. Our results provide evidence that Cxcr4 signaling also regulates neurofibromin-dependent cognitive function.

## Introduction

Neurofibromatosis type 1 (NF1) is an autosomal dominant, neurogenetic disorder caused by loss of function mutations in the gene neurofibromin 1 (*NF1*). NF1 is a comparatively common single gene disorder estimated to affect 1 in 2,000-3,000 live births (Evans et al., 2010; Huson et al., 1989; Uusitalo et al., 2015). NF1 is associated with a wide range of symptoms involving multiple organ systems. Neurological features, including peripheral nerve tumors, brain tumors, and cognitive dysfunction are major clinical concerns (Cimino and Gutmann, 2018). Up to 80% of children with NF1 experience cognitive or behavioral deficits, which impact social and academic development (Hyman et al., 2005; Torres Nupan et al., 2017). Despite increased attention on the role of cognitive function in quality of life for patients with NF1 (Krab et al., 2009; Varni et al., 2019), the molecular basis of NF1-associated cognitive dysfunction remains unclear (Miller and Halloran, 2022).

*NF1* encodes neurofibromin, a multifunctional GTPase-activating protein (GAP) that inhibits Ras signaling (Basu et al., 1992; Dasgupta et al., 2005; DeClue et al., 1991; Endo et al., 2013; Johannessen et al., 2005; Marchuk et al., 1991)and activates cAMP signaling (Anastasaki and Gutmann, 2014; Guo et al., 1997; Hannan et al., 2006). cAMP is a second messenger regulated by G protein-coupled receptor (GPCR) modulation of adenylyl cyclase (AC) activity. Dysregulated GPCR signaling is seen in neurofibromin-deficient human and mouse tissue, including increased CXCR4 chemokine receptor signaling associated with tumorigenesis (Karaosmanoglu et al., 2018; Mo et al., 2013; Warrington et al., 2007) and reduced dopamine signaling associated with cognitive function (Brown et al., 2011, 2010a; Diggs-Andrews et al., 2013). Ras and cAMP signaling also have been implicated in the regulation of neurofibromin-dependent cognitive function. Studies in both fruit flies and zebrafish show that loss of neurofibromin results in learning deficits that are rescued by enhancing the cAMP signaling pathway, whereas memory deficits are rescued by inhibiting the Ras signaling pathway (Ho et al., 2007; Wolman et al., 2014). Therefore, the role of neurofibromin might differ in neural circuits controlling distinct cognitive functions. The mechanism by which neurofibromin regulates cAMP signaling is unclear, as Ras-independent (Brown et al., 2012, 2010b; Guo et al., 1997), Ras-dependent (Anastasaki and Gutmann, 2014; Walker et al., 2006), and both Ras-independent and dependent pathways (Hannan et al., 2006) have been proposed.

Translation of research from animal models into treatments for NF1-associated cognitive dysfunction has seen limited success. Clinical trials for patients with NF1 using lovastatin or simvastatin to reduce Ras signaling have demonstrated limited effectiveness in improving learning or other measures of cognitive function (Bearden et al., 2016; Krab et al., 2008; Mainberger et al., 2013; Payne et al., 2016; Stivaros et al., 2018; van der Vaart et al., 2013). In a more recent clinical trial, regulation of Ras signaling through mitogen-activated protein kinase inhibitor (MEKi) treatment showed stable but not improved group-level performance on working memory and visual learning/memory tasks (Walsh et al., 2021). Identifying additional drug targets that improve learning in *Nf1* mutant animals and characterizing their mechanisms of action will provide new opportunities to develop treatments for cognitive dysfunction in patients with NF1.

In this study, using a zebrafish model of NF1 (Shin et al., 2012), we took two approaches to gain insight into the molecular mechanisms by which neurofibromin deficiency leads to behavioral and cognitive deficits. The first, a small-molecule drug screen to identify regulators of habituation learning, revealed that pharmacological inhibition of Cxcr4 chemokine receptor signaling with AMD3100 (Plerixafor) improves habituation deficits in *nf1* mutants. The second, RNAseq analysis to identify changes in gene expression with loss of neurofibromin, showed that *cxcr4b* is upregulated in *nf1* mutants. Cxcr4 signaling is known to activate Ras-Raf-MEK-ERK signaling and inhibit cAMP-PKA signaling (Busillo and Benovic, 2007). Therefore, we further investigated the effects of pharmacological inhibition of Cxcr4 on these signaling pathways. Using a whole-brain mitogen-activated protein kinase mapping (MAP-mapping) approach, we demonstrated that *nf1* mutant larvae have region-specific increases and decreases in phosphorylated-ERK (pERK). However, Plerixafor treatment produced limited changes in brain pERK levels in wild-type or *nf1* mutant larvae. In contrast, using an enzyme-linked immunosorbent assay (ELISA) and whole-brain immunolabeling, we show that *nf1* mutant larvae have decreased whole-body cAMP and whole-brain phosphorylated-PKA-substrate (pPKA-substrate) levels, which are increased by Plerixafor treatment in *nf1* heterozygous mutant larvae. Overall, we have identified Cxcr4 signaling as a regulator of neurofibromin-dependent habituation learning and modulator of cAMP-PKA signaling in *nf1* mutant larval zebrafish.

## Results

### Small-molecule screen reveals regulators of acoustically evoked behaviors in nf1 mutant larvae

A zebrafish model of NF1 harboring null alleles in *NF1* orthologs *nf1a* and *nf1b* was previously developed using a zinc finger nuclease strategy (Shin et al., 2012). We used these *nf1* mutants in an unbiased small-molecule screen to identify potential drug targets that regulate neurofibromin-dependent behaviors. Previous work showed that zebrafish *nf1* mutants have defects in prepulse inhibition (PPI) and habituation of the acoustic startle response, two measures of sensory filtering that demonstrate the ability of zebrafish larvae to adapt their behavior based on previous sensory experience (Randlett et al., 2019; Shin et al., 2012; Wolman et al., 2014). We measured the effects of pharmacological manipulation of cellular signaling on PPI and habituation learning in *nf1* mutant larvae.

In larval zebrafish, acoustic stimuli evoke robust startle responses consisting of ‘C-shape’ bends with distinct kinematic properties (Burgess and Granato, 2007a; Jain et al., 2018; Kimmel et al., 1974). Initiation of a short-latency C-bend (SLC) drives rapid escape behavior (Burgess and Granato, 2007a; Hecker et al., 2020; Jain et al., 2018; Marsden et al., 2018; Wolman et al., 2015). Our behavioral assay was designed to measure baseline startle initiation, PPI, and habituation of the zebrafish SLC response in *nf1* mutant larvae. A total of 60 acoustic stimuli were delivered with varying intensities and interstimulus intervals (ISIs) (Fig. 1A). Motor responses of individual larvae at 5 days post-fertilization (dpf) were recorded with a high-speed camera and response kinematics were analyzed using automated tracking software. The low-intensity stimulus was designed to evoke SLCs in response to approximately 25% of stimuli (Supp. Fig. 1A). The high-intensity stimulus was designed to evoke SLCs in response to approximately 75% of stimuli (Supp. Fig. 1A). PPI was assessed by comparing SLC initiation in response to individual high-intensity stimuli versus high-intensity stimuli that were preceded by low-intensity stimuli (Fig. 1A). Habituation was assessed by comparing SLC initiation in response to high-intensity stimuli separated by 1.5 second ISI instead of 20 second ISI (Fig. 1A).

**Figure 1.**
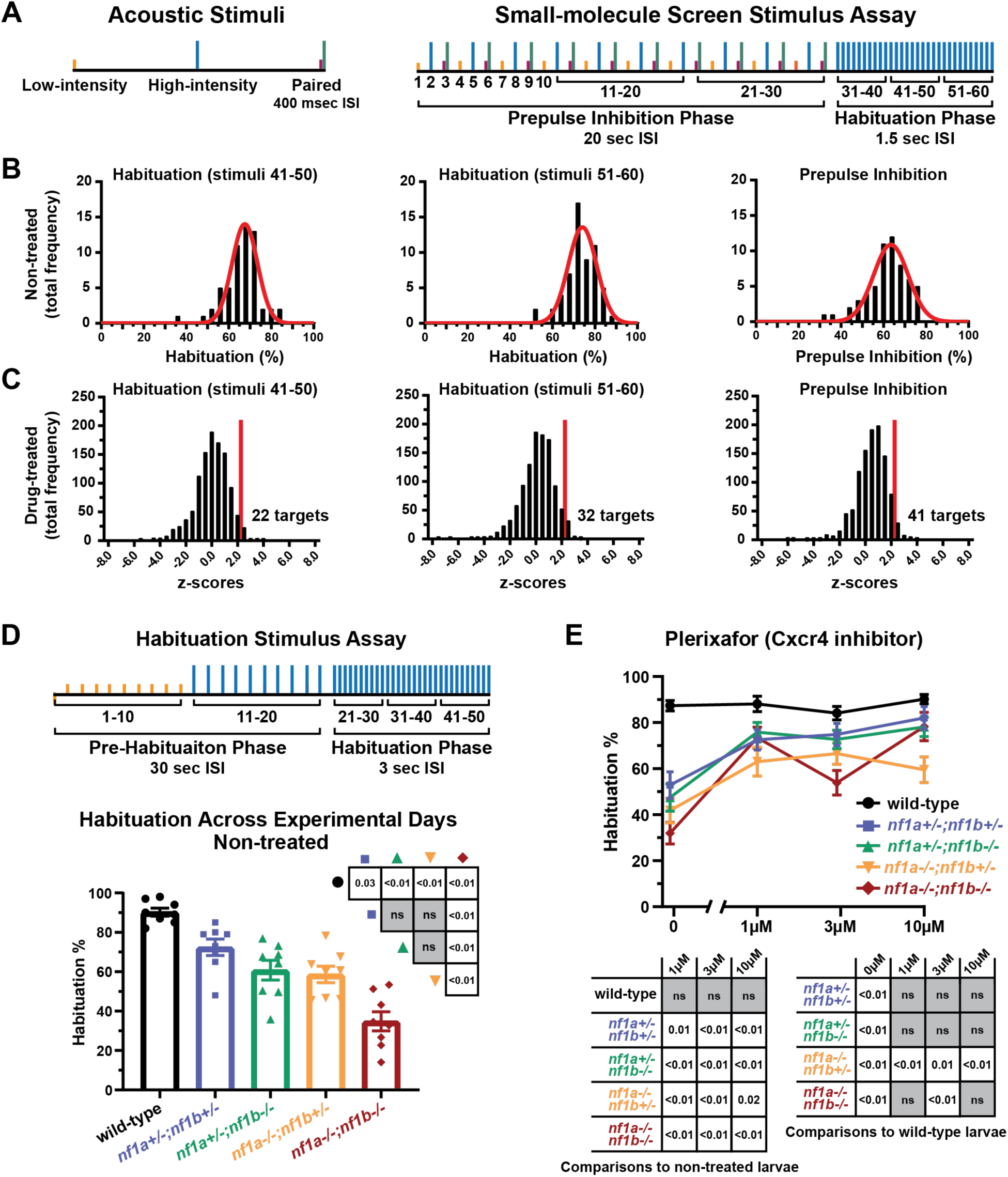
Small-molecule screen reveals Cxcr4 signaling regulates habituation in *nf1* mutant larvae. **(A)** Depiction of the stimulus assay used in combination with small-molecule screen. The low-intensity stimulus was designed to elicit 25% SLC responses. The high-intensity stimulus was designed to elicit 75% SLC responses. Paired low and high intensity stimuli were used during the prepulse inhibition (PPI) phase. The interstimulus interval (ISI) was reduced from 20 seconds to 1.5 seconds during the habituation phase. **(B)** Histograms demonstrating the frequency distribution of habituation and PPI percentage averaged across experimental days in 5 dpf non-treated *nf1* mutants. Each average was recorded from groups of 32 *nf1* mutant larvae. A nonlinear regression was fit to each distribution. The distribution average and standard deviation was used to calculate Z-scores to determine compounds that significantly affect behavioral measures. **(C)** Histograms demonstrating the frequency distribution of Z-scores calculated for effects of individual compounds. Each average was recorded from groups of 32 treated *nf1* mutant larvae. A Z-score threshold (2.326) was set representing the one-sided 99% confidence interval to identify compounds that improved habituation or PPI. **(D)** (Top) Depiction of the modified stimulus assay used to measure habituation. (Bottom) Habituation ± SEM in 5 dpf wild-type and *nf1* mutant larvae. Data points represent the average habituation from all larvae tested within each genotype on each experimental day (n=8). In total, 148-213 larvae were tested per genotype. One-way ANOVA showed a statistically significant difference between genotypes (F(4,35) = 23.62, p<0.001). Tukey’s adjusted p-values were used for comparisons between genotypes. **(E)** Habituation ± SEM for 5 dpf wild-type and *nf1* mutant larvae treated with Plerixafor. Data points represent average habituation from all larvae tested within each genotype at each treatment dose (sample sizes: WT control n=61, 1µM n=43, 3µM n=53, 10µM n=40; *nf1a^+/-^;nf1b^+/-^* control n=38, 1µM n=31, 3µM n=42, 10µM n=19; *nf1a^+/-^;nf1b^-/-^* control n=38, 1µM n=34, 3µM n=38, 10µM n=17; *nf1a^-/-^;nf1b^+/-^* control n=41, 1µM n=34, 3µM n=37, 10µM n=31; *nf1a^-/-^;nf1b^-/-^* control n=50, 1µM n=42, 3µM n=34, 10µM n=29). Two-way ANOVA showed statistically significant differences between treatment groups (F(3,732) = 31.11, p<0.001), as well as an interaction between the genotype and treatment factors F(12,732) = 4.017, p<0.001. Dunnet’s adjusted p-values (below graphs) were used for comparing non-treated larvae within each genotype and comparisons to wild-type larvae within each treatment dose.

Pharmacological screening of 1,134 small-molecule compounds with known biological targets was completed over the course of 60 experimental days. Pair mating *nf1a^+/-^*;*nf1b^-/-^* and *nf1a^-/-^*;*nf1b^+/-^* adults (Shin et al., 2012) yielded a mix of *nf1* mutant embryos with predicted equal ratios of *nf1a^+/-^;nf1b^+/-^*, *nf1a^+/-^;nf1b^-/-^*, *nf1a^-/-^;nf1b^+/-^*, and *nf1a^-/-^;nf1b^-/-^*mutant alleles. To determine baseline behavioral responsiveness in non-treated *nf1* mutant larvae, we quantified the baseline initiation of SLC responses and total motor reactions to low-intensity and high-intensity stimuli (Supp. Fig. 1A-B). To examine neurofibromin-dependent behaviors in non-treated *nf1* mutant larvae, we quantified habituation to stimuli 41-50, habituation to stimuli 51-60, and PPI (Fig. 1B). Data from two groups of 32 non-treated *nf1* mutant larvae were averaged on each experimental day and used as comparison to identify compounds that significantly regulated acoustically evoked behaviors.

To test the specific behavioral effects of compounds, we first analyzed drugs that altered baseline initiation of total reactions to low or high-intensity stimuli in *nf1* mutant larvae. 49 compounds increased baseline initiation of total reactions to low-intensity stimuli (Supp. Fig 1C; Supplementary Table 1), and 87 compounds decreased baseline initiation of total turns to high-intensity stimuli (Supp. Fig. 1C, Supplementary Table 1). We then performed enrichment analysis to identify classes of compounds with similar bioactivity using biological targets defined by the compound library manufacturer (Selleck). Compounds that increased baseline initiation of total reactions to low-intensity stimuli were enriched for retinoid receptor targets (Table 1). Compounds that decreased baseline initiation of total turns to high-intensity stimuli were enriched for estrogen and progestogen receptor, calcium channel, and 5-alpha reductase targets (Table 1). Although biologically interesting, we did not pursue compounds that altered baseline initiation of total turns in *nf1* mutant larvae and instead focused on compounds that specifically regulated habitation or PPI. 22 compounds improved habituation to stimuli 41-50 (Fig. 1C; Supplementary Table 1) and 32 compounds improved habituation to stimuli 51-60 (Fig. 1C; Supplementary Table 1). Compounds that improved habituation in *nf1* mutant larvae were enriched for adrenergic and 5-HT receptor targets (Table 1). Additionally, 41 compounds improved PPI (Fig. 1C; Supplementary Table 1). Compounds that improved PPI were enriched for 5-HT receptor, phosphodiesterase inhibitor, and topoisomerase inhibitor targets (Table 1).

**Table 1.** Enrichment analysis for Biological Targets that regulate acoustically evoked behaviors in *nf1* mutant larvae. Lists of biological targets enriched for target compounds that increase initiation of total reactions to low-intensity stimuli, decrease initiation of total reactions to high-intensity stimuli, increase habituation, and increase PPI in *nf1* mutants. Enrichment was determined by Fisher’s Exact test using a significance level of 0.05.

**Biological Targets Enriched for Increasing Total Reactions to Low-intensity Stimuli**
| Enriched Biological Target | Fisher's Exact p-value |
| --- | --- |
| Retinoid Receptor | 0.0104 |

**Biological Targets Enriched for Decreasing Total Reactions to High-intensity stimuli**
| Enriched Biological Target | Fisher's Exact p-value |
| --- | --- |
| 5-alpha Reductase | 0.0058 |
| Estrogen and progestogen Receptor | 0.0060 |
| Calcium Channel | 0.0430 |

**Biological Targets Enriched for Increasing Habituation Learning**
| Enriched Biological Target | Fisher's Exact p-value |
| --- | --- |
| Adrenergic (Habituation stimuli 51-60) | 0.0081 |
| 5-HT receptor (Habituation stimuli 51-60) | 0.0132 |
| 5-HT receptor (Habituation stimuli 41-50) | 0.0245 |

**Biological Targets Enriched for Increasing Prepulse Inhibition**
| Enriched Biological Target | Fisher's Exact p-value |
| --- | --- |
| 5-HT Receptor | 0.0063 |
| Phosphodiesterase Inhibitors | 0.0166 |
| Topoisomerase Inhibitors | 0.0393 |

Many of the compounds that improved habituation and/or PPI also decreased baseline initiation of total reactions to high-intensity stimuli (Supplementary Table 1). Of the drugs that specifically improved habituation and/or PPI without affecting baseline, we focused on the CXCR4 inhibitor Plerixafor. CXCR4 is a G_i_ protein-coupled chemokine receptor and its inhibition has been proposed as a treatment for optic glioma and malignant peripheral nerve sheath tumors (MPNSTs) in NF1 patients (Mo et al., 2013; Warrington et al., 2007). However, the effects of CXCR4 signaling on NF1-associated cognitive dysfunction have not been studied. Therefore, we further characterized the effects of pharmacological inhibition of Cxcr4 on habituation in wild-type and *nf1* mutant larvae.

### Cxcr4 signaling regulates habituation in nf1 mutant larvae

Previous studies of habituation of the SLC response in *nf1* mutant larvae identified deficits only in *nf1* double homozygous mutants (*nf1a^-/-^;nf1b^-/-^*) (Shin et al., 2012; Wolman et al., 2014). Because the human disease is autosomal dominant and manifests in heterozygous individuals, we established an assay sensitive to habituation deficits in heterozygous *nf1* mutants. We modified our original behavioral assay by removing the PPI phase and increasing the interstimulus interval (ISI) during the pre-habituation (30 seconds) and the habituation (3 seconds) phases (Fig. 1D). We tested non-treated wild-type and *nf1* mutant larvae in this modified assay, obtaining an average habituation value from groups of larvae from a total of 8 experimental days (Fig. 1D). Larvae were genotyped *post hoc* for *nf1a* and *nf1b* mutant alleles (Shin et al., 2012), and in total 148-213 larvae were tested per genotype. One-way ANOVA showed a statistically significant difference between genotypes (F(4,35) = 23.62, p<0.001). Adjusting for multiple comparisons between genotypes, habituation was significantly higher in wild-type larvae compared to each of the *nf1* mutant groups. Additionally, habituation was significantly lower in *nf1a^-/-^;nf1b^-/-^*mutant larvae compared to each of the *nf1* heterozygous (*nf1a^+/-^;nf1b^+/-^*, *nf1a^+/-^;nf1b^-/-^*, *nf1a^-/-^;nf1b^+/-^*) mutants (Fig. 1D). Using this modified behavioral assay, we were then able to test if pharmacological inhibition of Cxcr4 could improve habituation in both *nf1* double homozygous and *nf1* heterozygous mutant larvae.

We tested the effects of Cxcr4 inhibition on habituation by treating larvae with Plerixafor and Plerixafor 8HCl, two separate formulations of the Cxcr4 inhibitor AMD3100. Using both formulations (Fig. 1E; Supp. Fig. 1D), we found significant differences between groups. Plerixafor treatment (Fig. 1E) improved habituation in all *nf1* mutant genotypes without affecting wild-type habituation—restoring habituation to near wild-type levels in *nf1a^+/-^;nf1b^+/-^, nf1a^+/-^;nf1b^-/-^,* and *nf1a^-/-^;nf1b^-/-^* mutant larvae. Plerixafor 8HCl treatment (Supp. Fig. 1D) significantly improved habituation in *nf1a^-/-^;nf1b^-/-^* and *nf1a^+/-^;nf1b^-/-^* mutant larvae. Overall, these results support the conclusion that Cxcr4 signaling regulates habituation in *nf1* mutant larvae.

### Inhibition of Cxcr4 does not affect general motor performance at doses that improve habituation

To test the behavioral specificity of Cxcr4 inhibition at different treatment doses, we analyzed its effect on general motor performance by quantifying initiation of total turns to pre-habituation high-intensity stimuli. At the 3µM dose for Plerixafor and 300nM dose for Plerixafor 8HCl, treatment did not significantly affect baseline initiation of total turns to high-intensity stimuli in wild-type larvae (Fig. 2A; Supp. Fig. 1E). Therefore, we selected the 3µM dose of Plerixafor as a treatment that selectivity improves habituation in *nf1* mutant larvae (Fig. 1E) without affecting baseline initiation of turns (Fig. 2A). All subsequent experiments were conducted comparing non-treated and 3µM Plerixafor-treated larvae.

**Figure 2.**
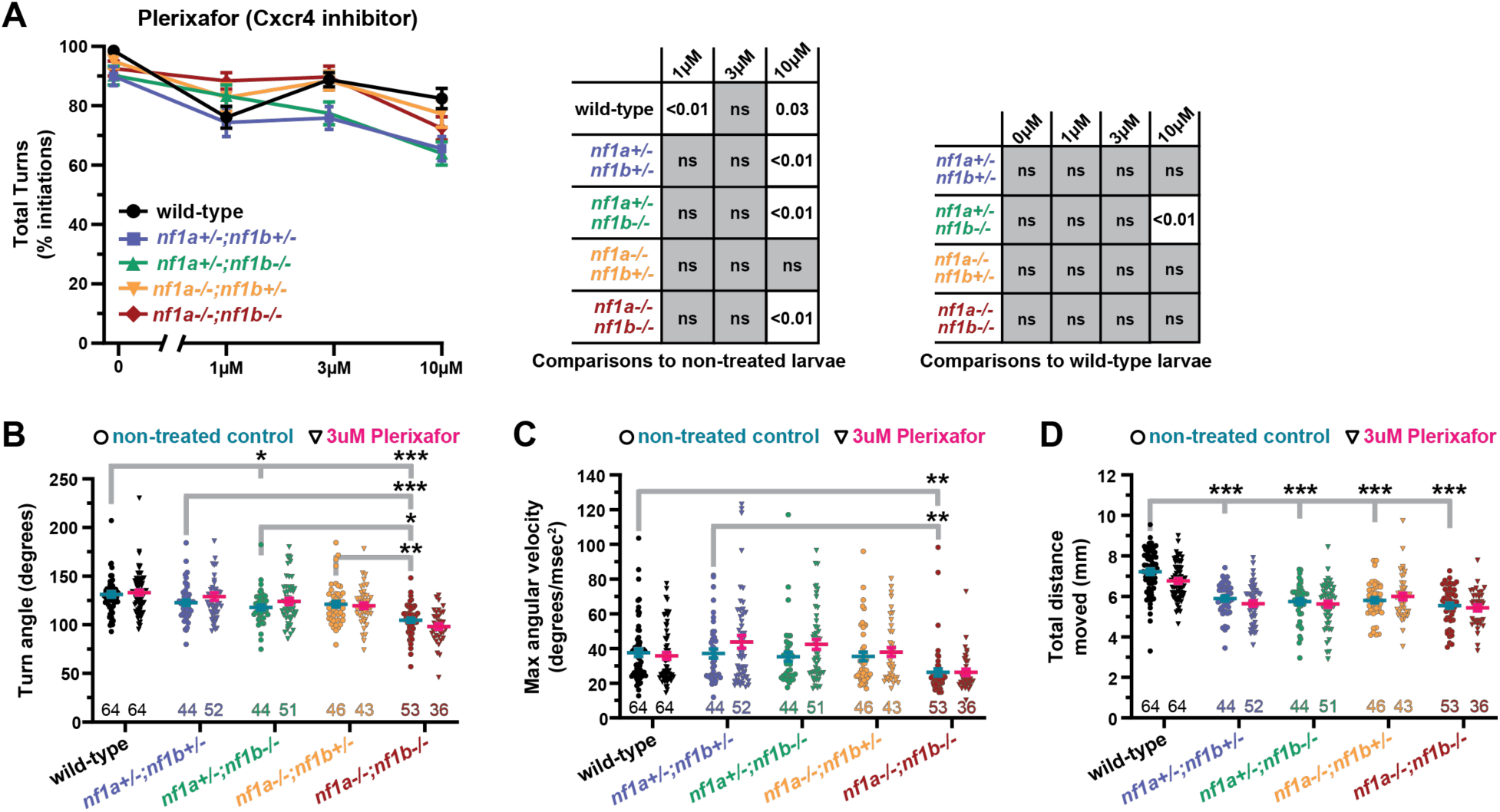
Inhibition of Cxcr4 does not affect general motor performance at doses that improve habituation. **(A)** Percent initiation of total reactions ± SEM in 5 dpf wild-type and *nf1* mutant larvae treated with Plerixafor. Data points represent average initiation percentage from all larvae tested within each genotype at each treatment dose (sample sizes: WT control n=63, 1µM n=64, 3µM n=63, 10µM n=62; *nf1a^+/-^;nf1b^+/-^* control n=44, 1µM n=45, 3µM n=53, 10µM n=39; *nf1a^+/-^;nf1b^-/-^* control n=44, 1µM n=39, 3µM n=50, 10µM n=52; *nf1a^-/-^;nf1b^+/-^* control n=45, 1µM n=44, 3µM n=42, 10µM n=42; *nf1a^-/-^;nf1b^-/-^* control n=53, 1µM n=51, 3µM n=37, 10µM n=50). Two-way ANOVA showed statistically significant differences between treatment groups (F(3,414) = 11.27, p<0.001), as well as a statistically significant interaction between genotype and treatment factors (F(12,962) = 1.907, p=0.030). Dunnet’s adjusted p-values were used for comparisons between non-treated larvae within each genotype **(B-D)** Turn angle, maximum angular velocity, and total distance moved in response to baseline high-intensity stimuli ± SEM in 5 dpf wild-type and *nf1* mutant larvae treated with 3µM Plerixafor. Data points represent average values from individual larvae in response to 10 high-intensity stimuli. Sample size indicated inside graph. Tukey’s adjusted p-values were used for comparing genotypes. **(B)** Two-way ANOVA showed statistically significant differences in turn angle between genotypes (F(4,487) = 27.80, p<0.001) but not Plerixafor treatment (F(1,487) = 0.4121, p=0.521). **(C)** Two-way ANOVA showed statistically significant differences in maximum angular velocity between genotypes (F(4,487) = 8.309, p<0.001) but not Plerixafor treatment (F(1,487 = 3.100, p=0.079). (**D)** Two-way ANOVA showed statistically significant differences in total distance moved between genotypes (F(4,487) = 42.29, p<0.001) but not Plerixafor treatment (F(1,487 = 2.735, p=0.099). * p<0.05, **p<0.01, ***p<0.001.

In a previous report on *nf1* mutant larvae, *nf1a^-/-^;nf1b^-/-^* mutants displayed motor deficits in addition to their habituation learning deficits, with kinematically weaker SLC responses as measured by decreased head turn angle, maximum angular velocity, and total distance traveled (Shin et al., 2012). To test if Cxcr4 inhibition affects the kinematic properties of turning behavior, we analyzed turn kinematics in non-treated and 3µM Plerixafor-treated larvae (Fig. 2B-D). Consistent with the previous report (Shin et al., 2012), we observed kinematically weaker SLC responses for turn angle, maximum angular velocity, and total distance moved in *nf1a^-/-^;nf1b^-/-^* larvae. Additionally, we found a decrease in total distance moved for each of the *nf1* heterozygous mutant larvae and a decrease in turn angle in *nf1a^+/-^;nf1b^-/-^* larvae. However, these motor deficits were not improved by inhibiting Cxcr4. Therefore, these results support a conclusion that Cxcr4 inhibition specifically regulates habituation learning in *nf1* mutants without altering general motor performance.

### Gene expression analysis reveals cxcr4b is upregulated in nf1 double homozygous mutants

We used a second approach, RNAseq analysis, to identify additional or overlapping molecular targets influenced by loss of neurofibromin. RNA was isolated from whole larvae in wild-type and putative *nf1* double homozygous mutant larvae identified by hyperpigmentation phenotype See Materials and Methods). Differential gene expression analysis followed by KEGG pathway enrichment analysis identified multiple metabolic pathways enriched for genes upregulated in *nf1* double homozygous mutants (Table 2). Focusing on Cxcr4 signaling, differential expression revealed upregulation of *cxcr4b* at 5 dpf in *nf1* double homozygous mutants compared to wild-type larvae (fold change = 1.52, adjusted p-value < 0.001). The increased expression of *cxcr4b* in *nf1* double homozygous mutant larvae alongside the small molecule drug screen results lead us to investigate the relationship between Cxcr4 and neurofibromin signaling.

**Table 2.**
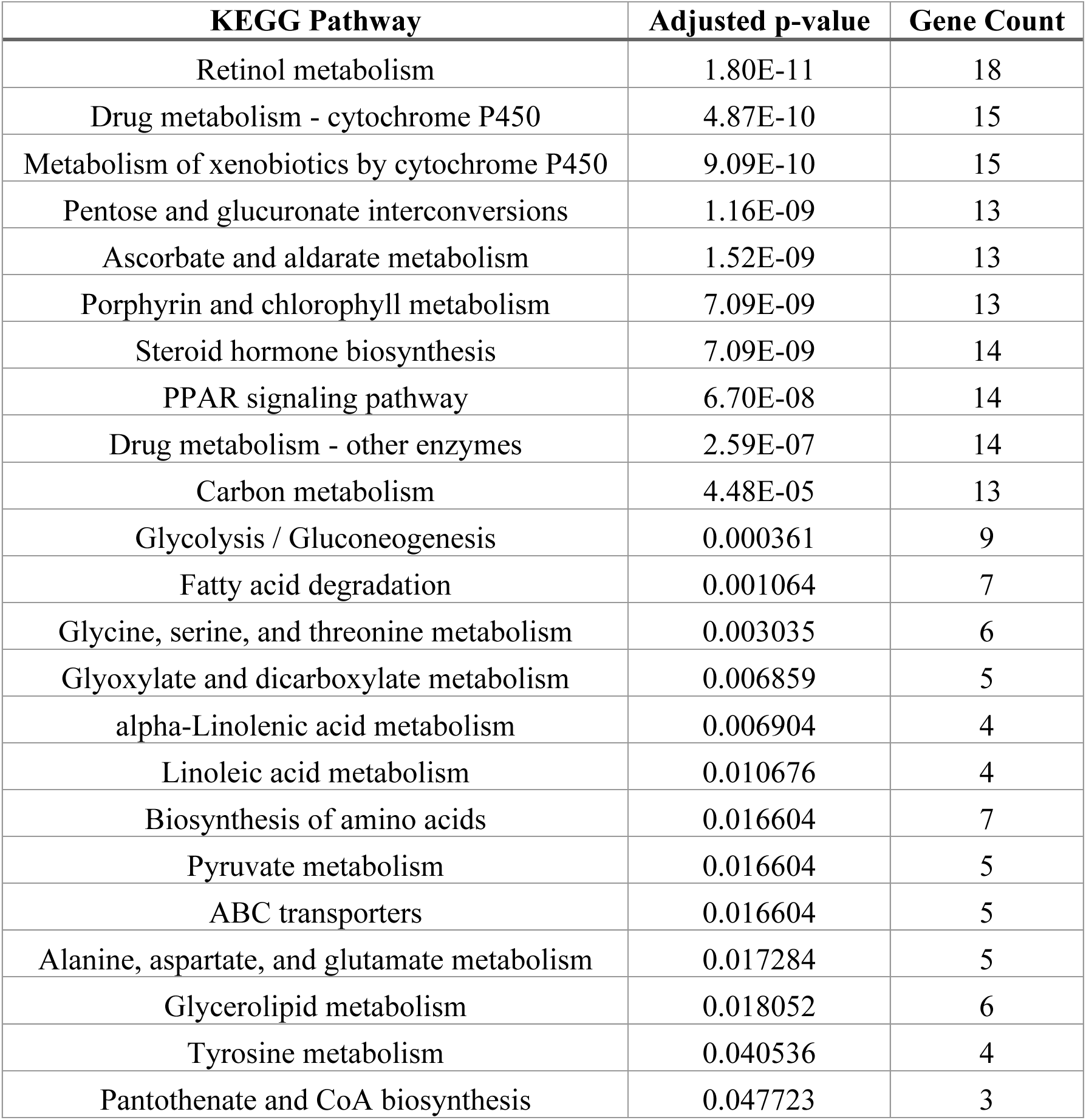
KEGG Pathway Analysis for Differentially Expressed Genes in *nf1* double homozygous mutants. Lists of KEGG pathways (Kyoto Encyclopedia of Genes and Genomes) with overrepresentation of differentially expressed genes from RNAseq analysis comparing 5-dfp *nf1* double homozygous mutants to wild-type larvae

### Cxcr4 signaling has limited effects on canonical Ras-Raf-MEK-ERK signaling in the brain

CXCR4 is a G_i_ protein-coupled chemokine receptor specific for the ligand stromal-derived-factor-1 (SDF-1 or CXCL12). Through Gα_i_ protein activation, the CXCR4 receptor is known to increase Ras signaling and inhibit cAMP signaling (Busillo and Benovic, 2007). Neurofibromin is also known to regulate these signaling pathways but in the opposing directions—by inhibiting Ras signaling and increasing cAMP signaling. Therefore, in neurofibromin-deficient larvae, we predicted Cxcr4 inhibition might decrease overactive Ras signaling and/or increase suppressed cAMP signaling.

We first tested where overactive Ras signaling was present in the brains of *nf1* mutant larvae. The canonical Ras signaling pathway involves a kinase cascade that phosphorylates mitogen-activated protein kinase (MAPK) also called extracellular signal-regulated kinase (ERK) (Lavoie et al., 2020). Phosphorylation of ERK can be detected and quantified in larval zebrafish brains using immunolabeling and the open-source Z-BRAIN reference atlas (Randlett et al., 2015). This technique, named MAP-mapping, quantifies differences in phosphorylation of ERK between experimental groups and maps those differences to brain regions of interest with cell-specific labels from reference lines. We fixed freely swimming larvae at 6 dpf, immunostained for total ERK (tERK) and phosphorylated ERK (pERK), and then processed images from wild-type, *nf1a^+/-^;nf1b^+/-^*, and *nf1a^-/-^;nf1b^-/-^*larvae. Comparing *nf1a^+/-^;nf1b^+/-^* larvae to wild-type (Fig. 3A) and *nf1a^-/-^;nf1b^-/-^*larvae to wild-type (Fig. 3B), we found region-specific increases and decreases in pERK. We focused on regions of interest associated with acoustic startle that displayed increased pERK (Table 3), an indicator that loss of neurofibromin may be contributing to overactivation of Ras in these regions. Acoustic startle responses in zebrafish are triggered by activation of reticulospinal Mauthner neurons in the hindbrain, which integrate acoustic and vibrational inputs from the statoacoustic ganglia and the lateral line and send output signals to motor neurons to induce muscle contraction (Eaton et al., 1977; Hecker et al., 2020; Liu and Fetcho, 1999; Wolman et al., 2015). Additionally, the octavolateralis nuclei and torus semicircularis relay auditory information to the thalamus. Some auditory information from the torus semicircularis and thalamus is also sent back to the hindbrain in zebrafish (Vanwalleghem et al., 2017) and could potentially provide top-down regulation of hindbrain activity to regulate acoustic startle habituation. Compared to wild type, *nf1a^+/-^;nf1b^+/-^*and *nf1a^-/-^;nf1b^-/-^* larvae showed increased pERK in the torus semicircularis and hindbrain, indicating that Ras signaling may be dysregulated in these regions. Therefore, we next asked if Cxcr4 inhibition decreases pERK in these brain regions.

**Figure 3.**
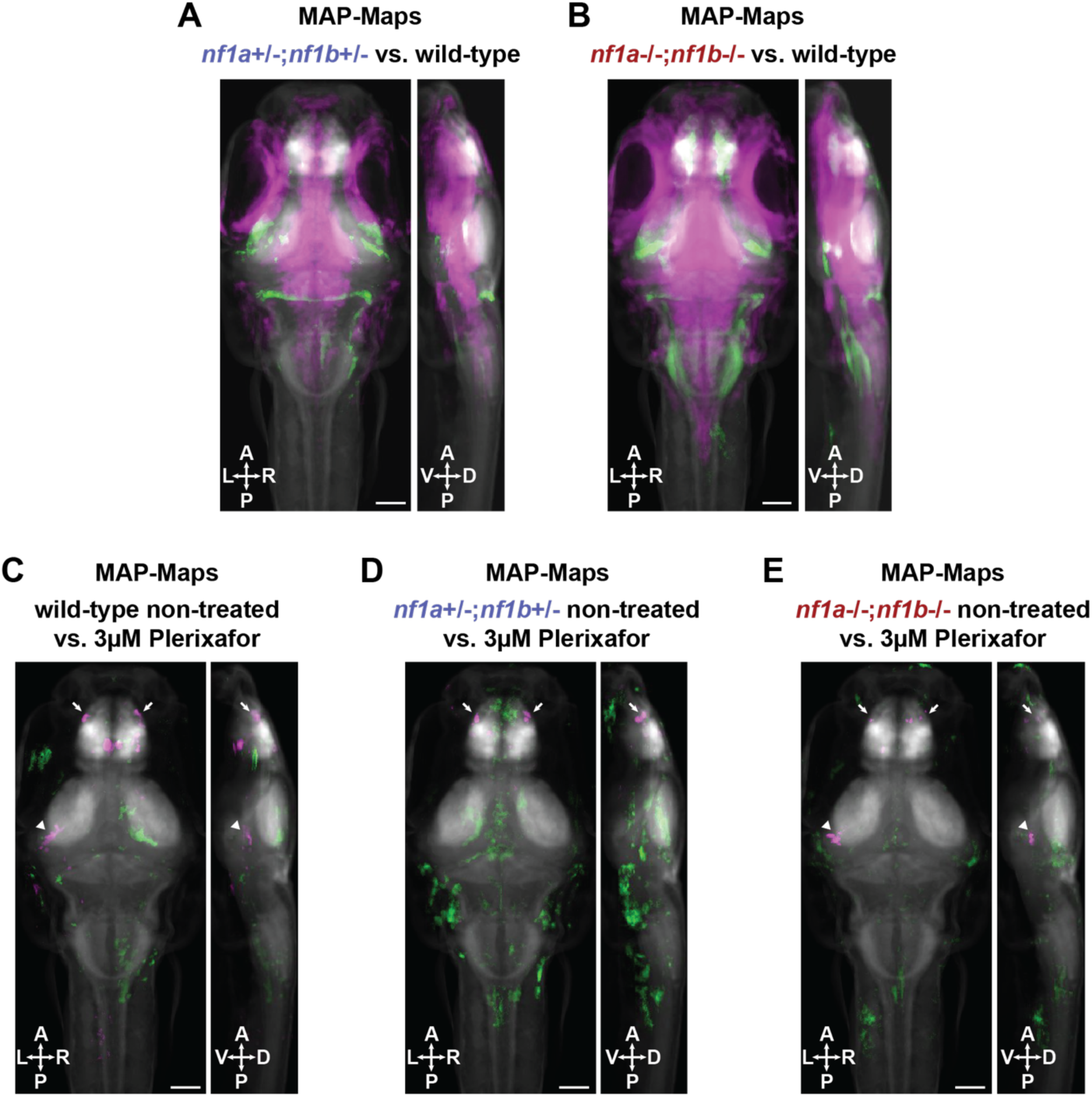
MAP-mapping of phosphorylated-ERK shows Cxcr4 inhibition has limited effects on canonical Ras signaling in the brain. **(A-B)** MAP-maps showing statistically significant median Z-score differences between non-treated **(A)** *nf1a^+/-^;nf1b^+/-^*versus wild-type larvae and **(B)** *nf1a^-/-^;nf1b^-/-^*versus wild-type larvae. Green signal represents increased pERK in *nf1* mutant larvae compared to wild-type. Magenta denotes decreased pERK in *nf1* mutant larvae compared to wild-type **(C-E)** MAP-maps showing statistically significant median Z-score difference between non-treated versus 3µM Plerixafor-treated **(C)** wild-type, **(D)** *nf1a^+/-^;nf1b^+/-^*, and **(E)** *nf1a^-/-^;nf1b^-/-^* larvae. Green signal represents increased pERK in Plerixafor-treated larvae. Magenta signal represents decreases pERK in Plerixafor-treated larvae. Arrowheads indicate region of interest torus semicircularis. Arrows indicate region of interest olfactory bulb. All larvae fixed and stained at 6 dpf. Z-scores calculated using Mann-Whitney U statistic with significance threshold set using a false discovery rate (FDR) of 0.00005. N = 18-20 larvae per treatment group. Scale bars represent 200µM.

**Table 3.** MAP-mapping region of interest analyses for *nf1* mutant versus wild-type larvae. Quantification of the mean Z-score difference between genotype groups in each region of interest is reported as Signal in ROI. The top 3 Z-BRAIN labels that show overlap with the Signal for each region of interest are reported. Reported signal values greater than 1 indicate relative enrichment.

| ROI analysis: increased pERK/tERK in *nf1a*+/-;*nf1b*+/- mutants vs. wild-type |  |  |  |  |  |  |  |
| --- | --- | --- | --- | --- | --- | --- | --- |
| Region of Interest | Signal in ROI | Cell-type Label 1 | Signal 1 | Cell-type Label 2 | Signal 2 | Cell-type Label 3 | Signal 3 |
| Torus Semicircularis | 426.6499 | Anti-5HT | 2.4756 | Anti-Gad67 | 2.1449 | Anti-GlyR | 2.1384 |
| Telencephalon (anterior forebrain) | 2.1172 | Anti-tERK | 4.6125 | Anti-5HT | 4.1013 | EtVmat2-GFP | 3.5844 |
| Diencephalon (posterior forebrain) | 35.4359 | Vglut2a-GFP | 2.4285 | Anti-tERK | 1.9462 | Anti-Zrf2 | 1.5595 |
| Mesencephalon (midbrain) | 193.4313 | Isl2bGal4-uasDendra | 30.6283 | Anti-Znp1 | 9.9624 | Anti-Zrf2 | 6.922 |
| Rhombencephalon (hindbrain) | 131.834 | Ptf1aGal4-uasKaede | 7.188 | Anti-Znp1 | 5.7143 | Anti-Zn1 | 5.6403 |

| ROI analysis: increased pERK/tERK in *nf1a*-/-;*nf1b*-/- mutants vs. wild-type |  |  |  |  |  |  |  |
| --- | --- | --- | --- | --- | --- | --- | --- |
| Region of Interest | Signal in ROI | Cell-type Label 1 | Signal 1 | Cell-type Label 2 | Signal 2 | Cell-type Label 3 | Signal 3 |
| Torus Semicircularis | 1290.011 | Anti-GlyR | 2.7734 | Anti-5HT | 2.7449 | Anti-Gad67 | 2.4702 |
| Telencephalon (anterior forebrain) | 1307.928 | Anti-Zrf2 | 6.5138 | Anti-tERK | 5.8422 | Anti-Zn1 | 4.4325 |
| Diencephalon (posterior forebrain) | 501.0858 | Anti-Zrf2 | 2.619 | Anti-tERK | 2.4348 | Vglut2a-GFP | 2.0668 |
| Mesencephalon (midbrain) | 344.4349 | Isl2bGal4-uasDendra | 15.7938 | Anti-Znp1 | 6.6898 | Anti-Zrf2 | 5.4395 |
| Rhombencephalon (hindbrain) | 635.2197 | Anti-GlyR | 9.0154 | Anti-Znp1 | 5.0124 | Anti-Zrf2 | 4.4444 |

To investigate whether Cxcr4 inhibition modulates Ras signaling in brain regions associated with acoustic startle, we performed MAP-mapping on wild-type (Fig. 3C), *nf1a^+/-^;nf1b^+/-^* (Fig. 3D), and *nf1a^-/-^;nf1b^-/-^* (Fig. 3E) larvae comparing non-treated and Plerixafor-treated groups (Table 4). Overall, Plerixafor treatment had limited effects on pERK in both wild-type and *nf1* mutant larvae. We did find decreased pERK in the torus semicircularis of Plerixafor treated wild-type and *nf1a^-/-^;nf1b^-/-^*larvae compared to non-treated larvae. However, these differences were unilateral and were not observed in *nf1a^+/-^;nf1b^+/-^* larvae (Fig. 3D). Additionally, Plerixafor treatment had very limited effects on pERK in the hindbrain (Table 4), where Mauthner neuron circuitry drives acoustic startle responses. Interestingly, the olfactory bulb showed decreased pERK in both wild-type and *nf1* mutant larvae treated with Plerixafor (Fig. 3C-E; Table 4). This is consistent with reports in larval zebrafish and adult mice showing *Cxcr4* mRNA is highly expressed in the olfactory bulb (Förster et al., 2017; Stumm et al., 2002). However, this result is unlikely to be related to regulation of acoustic startle responses. Taken together, these results support the conclusion that Cxcr4 signaling has a limited role in modulating canonical Ras-Raf-MEK-ERK signaling in the larval zebrafish brain.

**Table 4.** MAP-mapping region of interest analyses for Plerixafor-treated versus non-treated larvae. Quantification of the mean Z-score difference between treatment groups in each region of interest is reported as Signal in ROI. The top 3 Z-BRAIN labels that show overlap with the signal for each region of interest are reported. Reported signal values greater than 1 indicate relative enrichment.

| ROI analysis: decreased pERK/tERK in wild-type Plerixafor-treated vs. non-treated |  |  |  |  |  |  |  |
| --- | --- | --- | --- | --- | --- | --- | --- |
| Region of Interest | Signal in ROI | Cell-type Label 1 | Signal 1 | Cell-type Label 2 | Signal 2 | Cell-type Label 3 | Signal 3 |
| Torus Semicircularis | 97.6712 | Anti-5HT | 3.2096 | Anti-GlyR | 2.9163 | Anti-Gad67 | 2.5256 |
| Olfactory Bulb | 206.6698 | Vglut2a-GFP | 7.5826 | Elavl3-GCaMP5G | 4.3585 | Elavl3-H2BRFP | 4.1155 |
| Telencephalon (anterior forebrain) | 91.6901 | Anti-5HT | 5.874 | Anti-tERK | 5.866 | Anti-Zrf2 | 4.9997 |
| Diencephalon (posterior forebrain) | 49.4953 | Anti-Zrf2 | 1.9229 | Gad1b-GFP | 1.8527 | Anti-5HT | 1.4151 |
| Mesencephalon (midbrain) | 18.2628 | Anti-Znp1 | 3.5465 | Anti-GlyR | 3.5321 | Anti-5HT | 2.1924 |
| Rhombencephalon (hindbrain) | 2.0154 | Anti-GlyR | 2.775 | Elavl3-H2BRFP | 2.3555 | Vglut2a-GFP | 1.7154 |

| ROI analysis: decreased pERK/tERK in *nf1a+/-;nf1b+/-* Plerixafor-treated vs. non-treated |  |  |  |  |  |  |  |
| --- | --- | --- | --- | --- | --- | --- | --- |
| Region of Interest | Signal in ROI | Cell-type Label 1 | Signal 1 | Cell-type Label 2 | Signal 2 | Cell-type Label 3 | Signal 3 |
| Olfactory Bulb | 425.4882 | Vglut2a-GFP | 7.4568 | Elavl3-GCaMP5G | 4.0149 | Anti-Gad67 | 3.0747 |
| Telencephalon (anterior forebrain) | 63.0717 | Vglut2a-GFP | 5.8461 | Anti-tERK | 5.4957 | Anti-5HT | 5.39 |
| Diencephalon (posterior forebrain) | 2.0858 | Vglut2a-GFP | 4.3825 | Anti-tERK | 2.0935 | Anti-5HT | 1.8519 |
| Rhombencephalon (hindbrain) | 0.17789 | Anti-Zrf1(GFAP) | 3.1122 | Anti-GlyR | 2.298 | Anti-5HT | 1.57 |

| ROI analysis: decreased pERK/tERK in *nf1a-/-;nf1b-/-* Plerixafor-treated vs. non-treated |  |  |  |  |  |  |  |
| --- | --- | --- | --- | --- | --- | --- | --- |
| Region of Interest | Signal in ROI | Cell-type Label 1 | Signal 1 | Cell-type Label 2 | Signal 2 | Cell-type Label 3 | Signal 3 |
| Torus Semicircularis | 87.7413 | Anti-5HT | 3.1665 | Anti-GlyR | 3.038 | Anti-Gad67 | 2.9835 |
| Olfactory Bulb | 28.3954 | Vglut2a-GFP | 6.9219 | Anti-5HT | 2.6627 | Anti-Gad67 | 2.4144 |
| Telencephalon (anterior forebrain) | 10.7501 | Elavl3-H2BRFP | 4.1704 | Anti-5HT | 4.0196 | Vglut2a-GFP | 3.5166 |
| Diencephalon (posterior forebrain) | 1.0461 | Anti-Zrf2 | 4.6717 | Anti-tERK | 4.0909 | Anti-5HT | 2.7117 |
| Mesencephalon (midbrain) | 11.6061 | Anti-Znp1 | 4.7141 | Anti-GlyR | 4.2096 | Anti-Gad67 | 2.3958 |

### Cxcr4 signaling increases downregulated cAMP-PKA signaling in nf1 heterozygous mutants

Previous studies indicate that neurofibromin-deficient mice, fruit flies, and zebrafish have reduced cAMP levels (Brown et al., 2010b; Ho et al., 2007; Tong et al., 2002; Wolman et al., 2014). Consistent with these results, we found decreased cAMP levels in *nf1* mutants compared to wild-type larvae using an enzyme-linked immunosorbent assay (ELISA) to quantify cAMP (Fig. 4A). We collected tissue for ELISA from 5 dpf wild-type and *nf1* mutant larvae separated into *nf1* heterozygous mutant and *nf1* double homozygous mutant groups by hyperpigmentation phenotype. We found that cAMP levels were significantly lower in *nf1* heterozygous mutants compared to wild-type larvae and lower in *nf1* double homozygous mutants compared to both *nf1* heterozygous mutants and wild-type larvae (Fig. 4A). Next, we asked if Cxcr4 inhibition increased cAMP levels in *nf1* mutant larvae. We found that Plerixafor treatment increased cAMP levels in wild-type and *nf1* heterozygous mutant larvae but not *nf1* double homozygous mutant larvae (Fig. 4A), suggesting that CXCR4 inhibition of cAMP has an NF1-dependent component. Taken together, these results support the conclusion that *nf1* mutants have suppressed cAMP levels and that Cxcr4 inhibition raises cAMP levels in *nf1* heterozygous mutant larvae.

**Figure 4.**
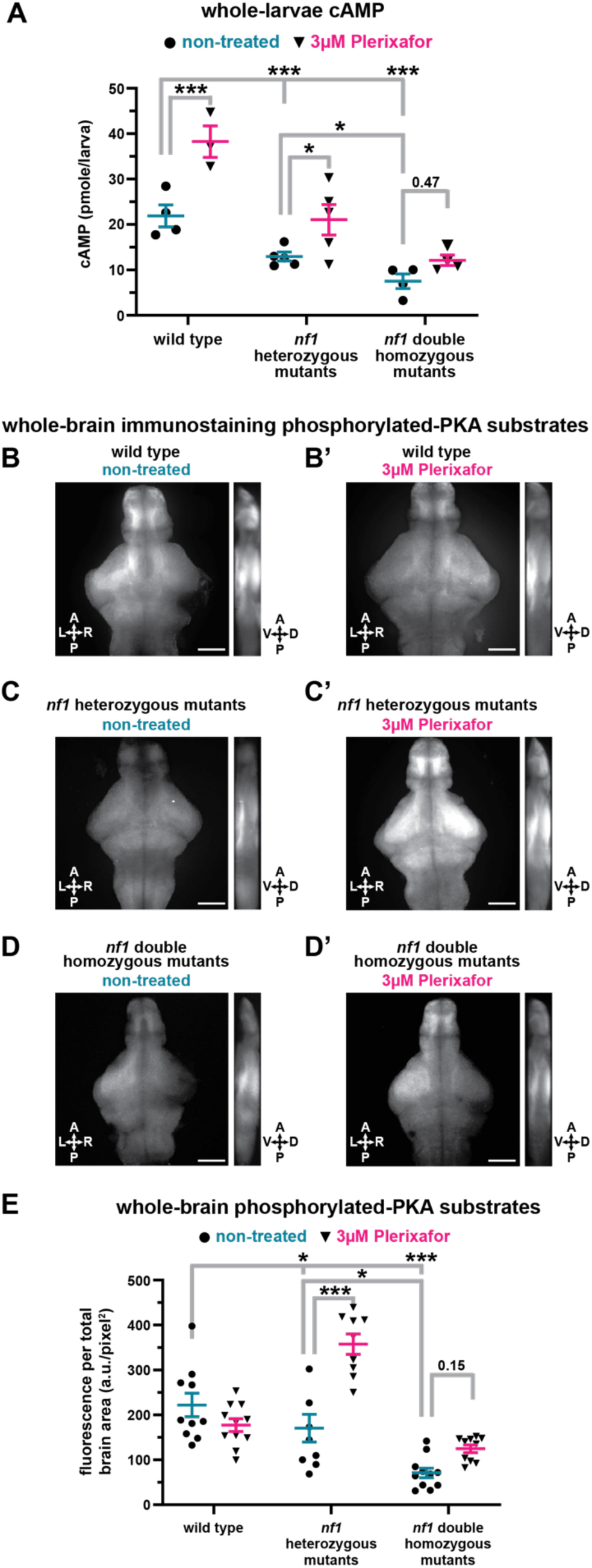
cAMP ELISA and phosphorylated-PKA substrate immunofluorescence show Cxcr4 inhibition increases downregulated cAMP-PKA signaling in *nf1* mutants. **(A)** cAMP levels ± SEM in 5 dpf wild-type and *nf1* mutant larvae treated with 3µM Plerixafor. Data points represent averages of 3 technical replicates (sample sizes: WT control n=4, treated n=3; *nf1* heterozygous mutant control n=5, treated n=5; *nf1* double homozygous mutant control n=4, treated n=4). Two-way ANOVA showed statistically significant differences between genotypes (F(2,19) = 33.79, p<0.001) and treatment groups (F(1,19) = 24.87, p<0.001). Tukey’s and Šídák’s adjusted p-values were used for multiple comparisons. **(B-D)** Representative phosphorylated-PKA substrate immunofluorescent images. Average intensity summation projections covering the whole brain. Scale bar represents 100µM. **(E)** phosphorylated-PKA substrate immunofluorescence. Values reported as arbitrary units of immunofluorescence per total brain area after subtracting background. Data points represent measurements from individual larvae (sample sizes: WT control n=10, treated n=11; *nf1* heterozygous mutant control n=8, treated n=9; *nf1* double homozygous mutant control n=11, treated n=10). Two-way showed statistically significant differences between genotypes F(2,54) = 35.72, p<0.001), treatments (F(1,54) = 16.41, p<0.001), and a statistically significant interaction between genotype and treatment (F(2,54) = 16.83, p<0.001). Tukey’s adjusted p-values were used for multiple comparisons. *p<0.05, **p<0.01, ***p<0.001

To regulate habituation of the SLC response, Cxcr4 signaling would need to modulate cAMP in neural circuits associated with acoustic startle. Therefore, we asked if Cxcr4 inhibition increases cAMP signaling in the brain. The cAMP effector protein kinase A (PKA) phosphorylates protein targets to regulate diverse cellular processes (Taskén and Aandahl, 2004). Using an antibody that recognizes substrates phosphorylated by PKA (pPKA-substrates) (Schmitt and Stork, 2002), we performed whole-brain immunolabeling on 5 dpf wild-type and *nf1* mutant larval brains. *nf1* heterozygous mutant and *nf1* double homozygous mutant larvae were separated into groups by hyperpigmentation phenotype. To enable the pPKA-substrate antibody to penetrate the brain, we dissected brain tissue from larvae and used a staining protocol that prevented us from registering images to the Z-BRAIN reference atlas. Instead, we created average fluorescence summation projections (Fig. 4B-D) to quantify pPKA-substrate fluorescence in whole brains (Fig. 4E). We found that *nf1* heterozygous mutants have significantly lower pPKA-substrate fluorescence than wild-type larvae and that *nf1* double homozygous mutants have significantly lower pPKA-substrate fluorescence than wild-type and *nf1* heterozygous mutants. With Plerixafor treatment, we found significantly increased pPKA-substrate fluorescence in *nf1* heterozygous mutant larvae only. Overall, these results support the conclusion that Cxcr4 signaling increases downregulated cAMP-PKA signaling caused by neurofibromin-deficiency.

## Discussion

Translation of research in animal models into treatments for NF1-associated cognitive dysfunction has not yet been successful, in part due to the complicated, multi-functional roles that neurofibromin plays in the distinct neural circuits and intracellular signaling cascades that regulate cognitive function. Here, we identify Cxcr4 chemokine receptor signaling as a regulator of habituation, a sensory filtering behavior and attention-based learning process similar to the cognitive behavior processes that are impaired in NF1 patients. While Cxcr4 signaling can regulate both the Ras and cAMP signaling pathways, we find that Cxcr4 predominantly affects cAMP-PKA signaling in *nf1* mutant larval zebrafish. Previous work showed that CXCR4 expression is elevated in malignant peripheral nerve sheath tumors (MPNSTs), and CXCR4 signaling was suggested as a target for treating neurofibromin-deficient tumorigenesis (Mo et al., 2013; Warrington et al., 2007). However, roles in NF1 cognitive dysfunction have not been explored. Our results provide evidence that Cxcr4 signaling may also regulate neurofibromin-dependent cognitive function and thus is a potential therapeutic target for NF1 cognitive deficits.

To identify molecular pathways that regulate neurofibromin-dependent behaviors, we chose to measure prepulse inhibition (PPI) and habituation of the SLC response in larval zebrafish. PPI and habituation learning deficits have previously been identified in neurofibromin-deficient mice and homozygous mutant zebrafish (Li et al., 2005; Randlett et al., 2019; Shin et al., 2012; Wolman et al., 2014). To develop a model that more closely resembles the heterozygous loss of function mutations observed in patients with NF1, we modified our behavioral assay to increase the interstimulus interval between acoustic stimuli during the habituation phase. This modification revealed behavioral deficits in *nf1* heterozygous mutants with at least one functioning allele of *nf1*, thus allowing us to better model the human disease.

Our behavioral drug screening experiments revealed Cxcr4 signaling as a modifier of behavioral deficits in *nf1* mutants. By testing multiple doses of two formulations of the Cxcr4 inhibitor Plerixafor in *nf1* mutants, we found robust increases in habituation at doses that did not affect baseline reactions to high-intensity stimuli, and that did not alter the kinematic properties of turning behavior. These findings indicate that Cxcr4 signaling specifically regulates habituation in *nf1* mutants without altering general motor performance.

NF1 and Cxcr4 signaling both affect the Ras-Raf-MEK-ERK and cAMP-PKA signaling pathways, but in opposite directions; NF1 decreases Ras activity and increases cAMP activity, while Cxcr4 increases Ras activity and suppresses cAMP activity. Thus, the finding that Cxcr4 inhibition rescues *nf1* mutant behavioral defects is perhaps not surprising, but prompts the question of the molecular mechanisms by which Cxcr4 signaling influences behavior in *nf1* mutants. Our finding that cAMP and PKA signaling were reduced in *nf1* mutant larvae, while increases in phosphorylation of ERK were limited and region-specific, suggests that cAMP signaling is the primary pathway influencing habituation behavior. These results are consistent with past work that investigated the molecular basis for learning and memory deficits in *nf1* mutant zebrafish and *Drosophila* and found that learning defects were rescued by manipulations that increase cAMP levels, while memory defects were rescued by inhibition of Ras signaling (Brown et al., 2010b; Ho et al., 2007; Tong et al., 2002; Wolman et al., 2014).

Our finding that Cxcr4 inhibition increased whole-body cAMP levels and brain pPKA-substrate immunofluorescence in nf1 heterozygous but not homozygous larvae suggests that a component of the Cxcr4 effect on cAMP and pPKA is neurofibromin-dependent. Cxcr4 and neurofibromin both influence multiple downstream signal transduction pathways with extensive possibilities for cross-talk and feedback within pathways. For example, cAMP-bound PKA regulatory subunits interact with Gαi and enhance Gαi signaling (Stefan et al., 2011). This represents a negative feedback mechanism in which high cAMP levels inhibit further adenylyl cyclase activity by potentiating Gi protein-coupled receptors such as Cxcr4. Therefore, the effect of Cxcr4 inhibition on cAMP-PKA signaling might be greater in *nf1* heterozygous larvae as a result of their higher baseline cAMP levels compared to *nf1* homozygous larvae. Surprisingly, we found that plerixafor treatment in wild-type larvae did not have a measurable effect on pPKA immunofluorescence in the brain. This finding may reflect a key role for neurofibromin in maintaining signaling pathway homeostasis. Under wild-type conditions and normal neurofibromin levels, the effects of plerixafor treatment on pPKA substrates may be transient and not detectable with the whole brain immunohistochemistry method. The complexity of the intertwined signaling pathways downstream from neurofibromin and Cxcr4 underscores the need for continued research into molecular mechanisms of NF1 disease.

It is interesting that total loss of neurofibromin in *nf1* double homozygous mutants does not lead to more global increases in pERK, as *nf1a* and *nf1b* are broadly expressed in the larval zebrafish brain (Padmanabhan et al., 2009). One possible explanation is that phosphorylation of ERK is also regulated by calcium signaling (Rosen et al., 1994; Thomas and Huganir, 2004). Dysregulation of intracellular calcium in neurons could potentially diminish the hyperactivation of Ras caused by loss of neurofibromin. Previous work in mice shows that loss of neurofibromin can lead to increased gamma aminobutyric acid (GABA) release in neural circuits (Cui et al., 2008; Shilyansky et al., 2010), which could inhibit calcium influx and thus inhibit the activation of Ras in affected brain regions.

Through MAP-mapping analysis, we did identify one brain region with increased pERK that could potentially regulate acoustically evoked startle habituation—the torus semicircularis (Fig. 3A; Table 3). In larval zebrafish, the torus semicircularis relays auditory information to the thalamus but also signals back to the hindbrain (Vanwalleghem et al., 2017), where it could potentially provide top-down regulation of hindbrain activity to regulate acoustic startle habituation. However, pERK in this region is not reduced in *nf1* heterozygous mutants by Cxcr4 inhibition (Fig. 3D; Table 4). Therefore, Cxcr4 signaling is unlikely to regulate habituation through modulation of the canonical Ras-Raf-MEK-ERK signaling pathway. This is consistent with a previous report that showed MEK inhibition with U0126 does not improve short-term habituation of the SLC response in larval zebrafish (Wolman et al., 2014). However, neurofibromin has also been shown to regulate cellular signaling through non-canonical Ras pathways (Anastasaki and Gutmann, 2014). Ras can activate atypical protein kinase C (PKCζ) to regulate cAMP homeostasis. If neurofibromin regulates cAMP through a Ras-PKCζ pathway in zebrafish larvae, it would not be detected with the pERK MAP-mapping technique used in this study. Therefore, we cannot rule out a potential role for non-canonical Ras signaling in the dysregulation of cAMP homeostasis we observe in *nf1* mutants.

### Other molecular targets that regulate neurofibromin-dependent behaviors

It is important to consider if the biological actions of the target compounds we identified are specific to loss of neurofibromin or if they affect behavior more generally. A small-molecule screen measuring short-term habituation of the SLC response was previously performed in wild-type larvae (Wolman et al., 2011). Target biological pathways identified in both our *nf1* mutant screen and the previous wild-type screen, including the 5-HT receptor pathway, may not be specific to loss of neurofibromin (Table 1; Wolman et al. 2011). Therefore, the effects of modulating serotonin signaling on habituation learning may not be specific to loss of neurofibromin. However, neurofibromin was shown to modulate 5-HT_6_ receptor constitutive activity in mouse neurons (Deraredj Nadim et al., 2016). Therefore, the relationship between neurofibromin and 5-HT receptor activity warrants further consideration. With respect to the compounds that target adrenergic receptors identified in our screen, the alpha-2 adrenergic receptor agonists are known to have sedative effects in humans and zebrafish (Giovannitti et al., 2015; Ruuskanen et al., 2005). Therefore, the effects of these compounds also may not be specific to habituation.

The compounds identified in our screen that improved PPI in *nf1* mutants were enriched for those targeting biological pathways that include 5-HT receptors, phosphodiesterase inhibitors, and topoisomerase inhibitors (Table 1). Modulation of 5-HT receptor signaling has previously been shown to regulate PPI in wild-type animals (Fletcher et al., 2001; Rigdon and Weatherspoon, 1992; Sipes and Geyer, 1994). Therefore, the effects of modulating serotonin signaling on PPI may not be specific to loss of neurofibromin. Phosphodiesterase (PDE) inhibitors increase cAMP signaling by preventing the degradation of cAMP. Interestingly, PDE inhibitors have previously been shown to improve habituation in *nf1* double homozygous mutants (Wolman et al., 2014) but their effects on PPI were unknown. Although compounds that improved habituation in our screen were not enriched for PDE inhibitors, we did identify Rolipram and Doxofylline as two PDE inhibitors that improve habituation in *nf1* mutants (Supplementary Table 1). Topoisomerases bind to and cut the DNA phosphate backbone and are used in medicine as antibiotics and chemotherapy agents. It is unclear how topoisomerase inhibitors might regulate PPI in *nf1* mutants. However, topoisomerase inhibitors can activate epigenetically silenced ubiquitin protein ligase alleles in neurons associated with Angelman syndrome (Huang et al., 2011), and mouse models for Angelman syndrome display deficits in acoustic startle and PPI (Huang et al., 2013). Further research will be required to understand how these effects might relate to neurofibromin.

Overall, we identify Cxcr4 chemokine receptor signaling as a regulator of habituation and modulator of cAMP-PKA signaling in *nf1* mutant larval zebrafish, suggesting that Cxcr4 signaling may regulate neurofibromin-dependent cognitive function in addition to its roles in tumorigenesis.

## Materials and Methods

### Fish maintenance and breeding

Zebrafish (*Danio rerio*) larvae used in this study were generated from crosses of wild-type (Tüpfel long fin background) adults or *nf1* mutant adults carrying a combination of *nf1a^Δ5^* (*nf1a ^-^*) and *nf1b^+10^* (*nf1b ^-^*) mutant alleles (Shin et al., 2012). Crossing *nf1a^+/-^*;*nf1b^-/-^* and *nf1a^-/-^*;*nf1b^+/-^*adults yielded embryos with *nf1a^+/-^*;*nf1b^+/-^*, *nf1a^+/-^*;*nf1b^-/-^*, *nf1a^-/-^*;*nf1b^+/-^*, or *nf1a^-/-^*;*nf1b^-/-^* mutant alleles. Genotyping of *nf1a^Δ5^* and *nf1b^+10^* alleles was performed as previously described (Shin et al., 2012). In indicated experiments, *nf1* mutant larvae were classified by the presence of a hyperpigmentation phenotype in *nf1a^-/-^*;*nf1b^-/-^* larvae. The resulting two groups of *nf1* mutant larvae are putative *nf1* heterozygous (*nf1a^+/-^*;*nf1b^+/-^*, *nf1a^+/-^*;*nf1b^-/-^*, *nf1a^-/-^*;*nf1b^+/-^*) and putative *nf1* double homozygous (*nf1a^-/-^*;*nf1b^-/-^*) mutants. This method of identification was validated by *post hoc* genotyping (100% positive predictive value and 93% negative predictive value for *nf1a^-/-^*;*nf1b^-/-^*, n=94)

Embryos and larvae were raised at 29°C in E3 medium (5 mM NaCl, 0.17 mM KCl, 0.33 mM CaCl2, 0.33 mM MgSO4, pH adjusted to 6.8–6.9 with NaHCO3) on a 14/10 h light/dark cycle. E3 medium was changed at 48 and 96 hours post fertilization (hpf). Before behavior testing, larvae were maintained on a white light box (800 μW/cm2) for at least 1 hour before being transferred to the testing arena. All experiments were conducted between 5 and 6 days post-fertilization (dpf).

### Behavioral Recording and Tracking

Behavioral responses to acoustically evoked startles were recorded as previously described (Burgess and Granato, 2007a; Wolman et al., 2011). Briefly, larvae housed individually in acrylic grids were exposed to acoustic/vibrational stimuli with a small vibration exciter (4810; Brüel and Kjaer, Norcross, GA) controlled by a digital–analog card (PCI-6221; National Instruments, Austin, TX). Larvae were illuminated from above with LED lights (MCWHL5 6500 K LED, powered by LEDD1B driver, Thorlabs) and below with infrared lights (IR Illuminator CM-IR200B, C&M Vision Technologies). Larval movement was recorded using MotionPro Y4 high-speed video cameras (Integrated Design Tools) with 50mm macro lenses (Sigma Corporation of America) at 1000 frames per second at either 512×512 or 1024×1024 pixel resolution.

Video images were analyzed for movement initiations and kinematics with the open-source Flote (v-2.0) software package as previously described (Burgess and Granato, 2007a, 2007b; Wolman et al., 2011). Data from individual larvae were matched to genotype information using a custom R script that is available upon request.

### Drug Screen Behavioral Assay

For the drug screen behavioral assay, *nf1* mutant larvae (*nf1a^+/-^*;*nf1b^+/-^*, *nf1a^+/-^*;*nf1b^-/-^*, *nf1a^-/-^*;*nf1b^+/-^*, or *nf1a^-/-^*;*nf1b^-/-^* mutant alleles, expected ratio 25% each) were tested at 5 dpf. A total of 60 acoustic stimuli were delivered with varying intensities and interstimulus intervals (ISIs) to assess baseline startle initiation, prepulse inhibition (PPI), and habituation (Fig. 1A). During the PPI phase (stimuli 1-30), a series of low-intensity, high-intensity, and paired low-high-intensity acoustic stimuli were delivered, separated by 20 second ISIs, with each stimulus repeated 10 times. The paired low-high-intensity stimuli were separated by 400ms. During the proceeding habituation phase (stimuli 31-60), 30 high-intensity acoustic stimuli were delivered, separated by 1.5 second ISIs.

Baseline startle initiation probability was assessed by measuring initiations (total turns or SLCs, separately) to both low and high-intensity stimuli during the PPI phase (stimuli 1, 2, 4, 5, 7, 8, 10, 11, 13, 14, 16, 17, 19, 20, 22, 23, 25, 26, 28, 29). The number of response initiations divided by the total analyzed was reported as the turn or SLC initiation percentage.

Habituation was assessed by calculating two ratios of SLC response initiation. The first was a ratio of SLC initiation in response to habituation phase stimuli 41-50 over SLC initiation in response to the first 10 high-intensity stimuli (PPI phase stimuli 2, 5, 8, 11, 14, 17, 20, 23, 26, 29). The second was a ratio of SLC initiation in response to stimuli 51-60 over SLC initiation in response to the first 10 high-intensity stimuli (PPI phase stimuli 2, 5, 8, 11, 14, 17, 20, 23, 26, 29). The habituation percentage is reported as 1 minus these ratios.

PPI was assessed by calculating the ratio of SLC initiations in response to paired low-high-intensity stimuli (PPI phase stimuli 3, 6, 9, 12, 15, 18, 21, 24, 27, 30) over SLC initiations in response to unpaired high-intensity stimuli (PPI phase stimuli 2, 5, 8, 11, 14, 17, 20, 23, 26, 29). The PPI percentage is reported as 1 minus this ratio.

### Habituation Behavioral Assay

The habituation behavioral assay was designed to be sensitive to more modest habituation deficits than the drug screen behavioral assay. Wild-type or *nf1* mutant larvae (genotyped *post hoc*) were tested in this assay at 5 dpf. A total of 50 acoustic stimuli were delivered with varying intensities and ISIs to assess baseline startle initiation and habituation (Fig. 2A). During the pre-habituation phase (stimuli 1-20), 10 low-intensity stimuli were delivered, followed by 10 high-intensity stimuli, all separated by 30 second ISIs. During the proceeding habituation phase (stimuli 21-50), 30 high-intensity stimuli were delivered, separated by 3 second ISIs.

Baseline startle initiation probability was assessed by measuring initiations (total turns or SLCs, separately) to both low and high-intensity stimuli during the pre-habituation phase (stimuli 1-20). The number of response initiations divided by the total analyzed was reported as the startle initiation percentage.

Habituation was assessed by calculating the ratio of SLC initiations in response to the final 10 high-intensity stimuli (habituation phase stimuli 41-50) over SLC initiations in response to the first 10 high-intensity stimuli (pre-habituation phase stimuli 11-20). The habituation percentage is reported as 1 minus this ratio.

### Pharmacology

For the small-molecule screen, 1,134 bioactive compounds from an FDA-approved small-molecule library (Selleck) were tested on *nf1* mutant larvae. Compounds were maintained at - 80°C dissolved in DMSO and were administered to larvae at a final concentration of 1µM or 10µM per compound (in 1% DMSO in E3 medium) for a treatment duration of 1 hour before behavioral testing. Each compound was administered to 32 larvae (mix of *nf1* mutant genotypes). 2 groups of 32 larvae (mix of *nf1* mutant genotypes) were left untreated on each experimental day. Behavioral testing was completed over the course of 60 experimental days.

For follow-up habituation experiments, two formulations of the Cxcr4 inhibitor AMD3100 (Plerixafor 8HCl and Plerixafor) were tested on wild-type and *nf1* mutant larvae (genotyped *post hoc*). Both formulations were dissolved in water and administered to larvae in a series of doses for a treatment duration of 1 hour before behavioral testing. For Plerixafor 8HCl, the final concentrations in E3 medium were 300nM, 1µM, and 3µM. For Plerixafor, the final concentrations in E3 medium were 1µM, 3µM, and 10µM. For remaining experiments (measures of ERK, cAMP, and PKA signaling), the 3µM Plerixafor dose was used as treatment for a duration of 1 hour.

### RNA sequencing and differential gene expression analysis

RNA was isolated from groups of 20 whole-larvae at 5 dpf using TRIzol Reagent (Invitrogen) following the manufacturer’s instructions. 8 biological replicates for wild-type (Tüpfel long fin background) and putative *nf1* double mutants were processed for RNA sequencing. RNA sequencing was carried out by the Biotechnology Center Gene Expression Center at the University of Wisconsin-Madison on an Illumina HiSeq 2500 platform. Transcriptomic data processing adhered to ENCODE Consortium guidelines and best practices for RNA-Seq (ENCODE Project Consortium, 2016). Raw strand-specific paired-end (2×150 bp) Illumina reads were adapter-trimmed using Skewer (v0.1.123; Jiang et al. 2014). Trimmed reads were aligned to the *Danio rerio GRCz11* reference genome (NCBI Assembly: GCA_000002035.4) using STAR (v2.5.3a; Dobin et al. 2013), a splice-aware aligner, with gene annotation from Ensembl release 102. Gene- and isoform-level expression quantification was performed with RSEM (v1.3.1; Li and Dewey 2011), which estimates transcript abundance using expectation-maximization. For downstream differential gene expression analysis, expected read counts from RSEM were rounded to the nearest integer and imported into edgeR (v3.28.0; Robinson, McCarthy, and Smyth 2010), run within the R statistical environment (v3.6.2) Library size normalization across samples was performed using the trimmed mean of M-values (TMM) method (Robinson and Oshlack 2010). Genes were retained for statistical testing if they exceeded a count-per-million (CPM) threshold in a minimum number of samples, defined by the number of biological replicates per condition, with a minimum read count of 10. This independent filtering step reduced noise and increased statistical power. Differential expression testing was conducted using negative binomial generalized linear models. Resulting *p*-values were adjusted for multiple testing using the Benjamini–Hochberg false discovery rate (FDR) procedure, with a significance threshold of 5% (Reiner et al., 2003). The validity of the multiple testing correction was assessed by visual inspection of the distribution of unadjusted *p*-values. KEGG pathway analysis was performed as described in Subhash and Kanduri (Subhash and Kanduri, 2016).

### Immunohistochemistry and confocal imaging

Immunostaining and whole-brain imaging for total-ERK (tERK) and phosphorylated-ERK (pERK) was performed as previously described (Randlett et al., 2015). Larvae at 6 dpf were fixed in 4% paraformaldehyde (diluted to 4% w/v in PBS from 16% w/v in 0.1 M phosphate buffer, 0.25% v/v Triton-X, pH 7.4) overnight at 4°C. The next day, larvae were incubated in bleaching solution (3% v/v hydrogen peroxide, 1% w/v KOH, in water) for 30 minutes to remove pigment. Larvae were then incubated in 150mM TrisHCl pH 9.0 for 15 minutes at 70°C. Larvae were permeabilized in 0.05% Trypsin-EDTA on ice for 45 minutes. Larvae were then incubated for 1 hour in block solution (1% w/v bovine serum albumin, 2% v/v normal goat serum, 0.25% v/v Triton-X, 1% v/v DMSO, in PBS, pH 7.4). Larvae were then incubated in primary antibodies at 1:300 in incubation buffer (IB; 1% w/v bovine serum albumin, 0.25% v/v Triton-X, 1% v/v DMSO, in PBS, pH 7.4) overnight at 4°C. The next day, larvae were incubated in fluorescently conjugated secondary antibodies at 1:500 in IB overnight at 4°C and stored in Vectashield mounting media (Vector Laboratories) until mounting for imaging. Primary antibodies included phospho-ERK antibody (Cell Signaling, #4370) and total-ERK antibody (Cell Signaling, #4696). Secondary antibodies included AlexaFluor488-conjugated goat anti-mouse IgG1 and AlexaFluor594-conjugated goat anti-rabbit IgG (ThermoFisher Scientific) Whole-brain image stacks were acquired on an Olympus Fluoview confocal laser scanning microscope (FV1000) using a 20x oil immersion objective and Fluoview software (FV10-ASW 4.2). Images were acquired at a x/y/z resolution of 0.795/0.795/2uM. The entire brain was captured by imaging in two sections and stitching images using the Pairwise Stitching plugin in FIJI. Pixel saturation was adjusted to 0.1% of the most intense pixels for each sample using a custom FIJI macro available upon request.

Whole-brain image stacks for total-ERK (tERK) and phosphorylated-ERK (pERK) were registered to the open-source Z-BRAIN atlas using the Jefferis lab graphical user interface for the Computational Morphometry Toolkit (CMTK) hosted on GitHub. Parameters used included: - awr 0102 -X 52 -C 8 -G 80 -R 3 -A ’--accuracy 0.4’ -W ‘--accuracy 1.6’. Images were downsampled and smoothed using the PrepareStacksForMAPMapping FIJI macro. MAP-maps were created using the MakeTheMAPMap MATLAB script. Z-BRAIN analysis of MAP-maps was generated using the ZbrainAnalysisOfMAPMaps. FIJI and MATLAB scripts available from Z-BRAIN hosted by the Zuse Institut Berlin.

Immunostaining and whole-brain imaging was performed on dissected brains for phosphorylated-PKA substrates (pPKA-substrates). Larvae at 5 dpf were fixed in 4% paraformaldehyde (diluted to 4% w/v in PBS from 16% w/v in 0.1 M phosphate buffer, 5% w/v sucrose, pH 7.4) for 1 hour at room temperature. Brains were dissected from the head using fine tungsten dissecting needles. Brains were permeabilized in collagenase (0.1% w/v in PBS) for 1 hour and blocked for 1 hour at room temperature in incubation buffer (IB; 0.2% w/v bovine serum albumin, 2% v/v normal goat serum, 0.8% v/v Triton-X, 1% v/v DMSO, in PBS, pH 7.4). Brains were incubated in primary antibodies at 1:300 in IB overnight at 4°C. Brains were incubated in fluorescently conjugated secondary antibodies at 1:500 in IB overnight at 4°C and stored in Vectashield mounting media (Vector Laboratories) until mounting for imaging. Primary antibodies included phospho-(Ser/Thr) PKA substrate (Cell Signaling, #9621) and total-ERK (Cell Signaling, #4696). Secondary antibodies included AlexaFluor488-conjugated goat anti-mouse IgG1 and AlexaFluor594-conjugated goat anti-rabbit IgG (ThermoFisher Scientific). Whole-brain image stacks were acquired on an Olympus Fluoview confocal laser scanning microscope (FV1000) using a 20x oil immersion objective and Fluoview software (FV10-ASW 4.2). Images were acquired at a x/y/z resolution of 0.795/0.795/2uM.

Whole-brain image stacks for pPKA-substrates were processed using the open-source software FIJI (v-1.53c) (Schindelin et al., 2012). Whole-brain pPKA-substrates fluorescence was measured by drawing a region of interest around the entire brain from Z-stack average intensity projections. Background fluorescence was measured by drawing a region of interest away from sample in the same projection. The total corrected fluorescence was quantified using the following calculation: Integrated Density – (Area of region of interest x Mean fluorescence of the background). Fluorescence was reported as total corrected fluorescence per total brain area.

### cAMP enzyme-linked immunosorbent assay (ELISA)

For cAMP quantification, zebrafish larvae at 5 dpf were snap-frozen in liquid nitrogen. Tissue from 15 larvae per group was lysed by homogenizing in 200 uL of 0.1 M HCl using a hand-held homogenizer. Samples were spun at 12,500 rpm for 15 minutes at 4°C. The supernatant was transferred to a new microcentrifuge tube and diluted 1:1 in 0.1 M HCl. cAMP concentration was quantified using a direct cAMP ELISA kit (Enzo Life Science, ADI900066) following the acetylated version of the manufacturer’s instructions. Three technical replicates were averaged for each sample reported.

### Statistical Analysis

Descriptive statistics (means, standard deviations, standard errors) and Z-scores were calculated in the software environments RStudio (v-1.3), GraphPad Prism (v-9.1.2, GraphPad Software Incorporated), and Matlab (v-9.6.0, R2019A, The MathWorks Inc.). Z-scores for the effects of individual compounds in the small-molecule screen were calculated with the following formula: Z-score = (Mean treated value – Mean non-treated control value) / (Standard deviation non-treated control value). Z-scores were identified as significant by setting a critical value based on the one-tailed 99% confidence interval. Enrichment analysis for target compounds was performed using Fisher’s exact test. Assumptions of normality were tested by Shapiro-Wilk’s test. One-way ANOVA, two-way ANOVA, and Tukey’s and Dunnet’s multiple comparisons testing were performed with a significance level of 0.05. Gene expression analysis was performed using R (v-3.6.2). Data are presented as means ± standard error of the mean (SEM). Sample sizes and statistical tests for specific experiments are identified in the figures and figure legends.

## Acknowledgements

The authors would like to thank Dr. Marc Wolman for technical advice on the small-molecule compound behavioral screen, Dr. Owen Randlett for technical advice on the MAP-mapping procedure, Dr. Changjiu Zhao for technical assistance with ELISA experiments, and Riley Kelley, Morgan Wetzel, Hollis Howe, Liza Rick, Hannah Olson, Jessa Snower, and Blanche Danecek for technical assistance, Dr. Michael Granato for the *nf1* mutant fish lines, and the University of Wisconsin-Madison Biotechnology Center: Gene Expression Center (RRID:SCR_017757) for RNA library preparation; DNA Sequencing Facility (RRID:SCR_017759) for sequencing; and Bioinformatics Resource Center (RRID:SCR_017799) for analysis services.

## Author Contributions

Conceptualization: A.H.M., M.C.H.; Validation: M.E.B.; Formal analysis: A.H.M., Y.Y., N.S., M.E.B.; Investigation: A.H.M., Y.Y., N.S., J.M.; Resources: M.C.H.; Writing – original draft preparation: A.H.M.; Writing – review and editing: A.H.M., M.E.B., M.C.H.; Visualization: A.H.M.; Supervision: M.C.H.; Funding acquisition: M.C.H.

## References

Anastasaki, C., Gutmann, D.H., 2014. Neuronal NF1/RAS regulation of cyclic AMP requires atypical PKC activation. Hum Mol Genet 23, 6712–6721. 10.1093/hmg/ddu389

Basu, T.N., Gutmann, D.H., Fletcher, J.A., Glover, T.W., Collins, F.S., Downward, J., 1992. Aberrant regulation of ras proteins in malignant tumour cells from type 1 neurofibromatosis patients. Nature 356, 713–715. 10.1038/356713a0

Bearden, C.E., Hellemann, G.S., Rosser, T., Montojo, C., Jonas, R., Enrique, N., Pacheco, L., Hussain, S.A., Wu, J.Y., Ho, J.S., McGough, J.J., Sugar, C.A., Silva, A.J., 2016. A randomized placebo-controlled lovastatin trial for neurobehavioral function in neurofibromatosis I. Ann Clin Transl Neurol 3, 266–279. 10.1002/acn3.288

Brown, J.A., Diggs-Andrews, K.A., Gianino, S.M., Gutmann, D.H., 2012. Neurofibromatosis-1 heterozygosity impairs CNS neuronal morphology in a cAMP/PKA/ROCK-dependent manner. Mol Cell Neurosci 49, 13–22. 10.1016/j.mcn.2011.08.008

Brown, J.A., Emnett, R.J., White, C.R., Yuede, C.M., Conyers, S.B., O’Malley, K.L., Wozniak, D.F., Gutmann, D.H., 2010a. Reduced striatal dopamine underlies the attention system dysfunction in neurofibromatosis-1 mutant mice. Hum Mol Genet 19, 4515–4528. 10.1093/hmg/ddq382

Brown, J.A., Gianino, S.M., Gutmann, D.H., 2010b. Defective cAMP generation underlies the sensitivity of CNS neurons to neurofibromatosis-1 heterozygosity. J Neurosci 30, 5579–5589. 10.1523/JNEUROSCI.3994-09.2010

Brown, J.A., Xu, J., Diggs-Andrews, K.A., Wozniak, D.F., Mach, R.H., Gutmann, D.H., 2011. PET imaging for attention deficit preclinical drug testing in neurofibromatosis-1 mice. Exp Neurol 232, 333–338. 10.1016/j.expneurol.2011.09.005

Burgess, H.A., Granato, M., 2007a. Sensorimotor gating in larval zebrafish. J Neurosci 27, 4984–4994. 10.1523/JNEUROSCI.0615-07.2007

Burgess, H.A., Granato, M., 2007b. Modulation of locomotor activity in larval zebrafish during light adaptation. Journal of Experimental Biology 210, 2526–2539. 10.1242/jeb.003939

Busillo, J.M., Benovic, J.L., 2007. Regulation of CXCR4 signaling. Biochim Biophys Acta 1768, 952–963. 10.1016/j.bbamem.2006.11.002

Cimino, P.J., Gutmann, D.H., 2018. Neurofibromatosis type 1. Handb Clin Neurol 148, 799–811. 10.1016/B978-0-444-64076-5.00051-X

Cui, Y., Costa, R.M., Murphy, G.G., Elgersma, Y., Zhu, Y., Gutmann, D.H., Parada, L.F., Mody, I., Silva, A.J., 2008. Neurofibromin regulation of ERK signaling modulates GABA release and learning. Cell 135, 549–560. 10.1016/j.cell.2008.09.060

Dasgupta, B., Yi, Y., Chen, D.Y., Weber, J.D., Gutmann, D.H., 2005. Proteomic analysis reveals hyperactivation of the mammalian target of rapamycin pathway in neurofibromatosis 1-associated human and mouse brain tumors. Cancer Res 65, 2755–2760. 10.1158/0008-5472.CAN-04-4058

DeClue, J.E., Cohen, B.D., Lowy, D.R., 1991. Identification and characterization of the neurofibromatosis type 1 protein product. Proc Natl Acad Sci U S A 88, 9914–9918. 10.1073/pnas.88.22.9914

Deraredj Nadim, W., Chaumont-Dubel, S., Madouri, F., Cobret, L., De Tauzia, M.-L., Zajdel, P., Bénédetti, H., Marin, P., Morisset-Lopez, S., 2016. Physical interaction between neurofibromin and serotonin 5-HT6 receptor promotes receptor constitutive activity. Proc Natl Acad Sci U S A 113, 12310–12315. 10.1073/pnas.1600914113

Diggs-Andrews, K.A., Tokuda, K., Izumi, Y., Zorumski, C.F., Wozniak, D.F., Gutmann, D.H., 2013. Dopamine deficiency underlies learning deficits in neurofibromatosis-1 mice. Ann Neurol 73, 309–315. 10.1002/ana.23793

Dobin, A., Davis, C.A., Schlesinger, F., Drenkow, J., Zaleski, C., Jha, S., Batut, P., Chaisson, M., Gingeras, T.R., 2013. STAR: ultrafast universal RNA-seq aligner. Bioinformatics 29, 15–21. 10.1093/bioinformatics/bts635

Eaton, R.C., Bombardieri, R.A., Meyer, D.L., 1977. The Mauthner-initiated startle response in teleost fish. J Exp Biol 66, 65–81. 10.1242/jeb.66.1.65

Endo, M., Yamamoto, H., Setsu, N., Kohashi, K., Takahashi, Y., Ishii, T., Iida, K., Matsumoto, Y., Hakozaki, M., Aoki, M., Iwasaki, H., Dobashi, Y., Nishiyama, K., Iwamoto, Y., Oda, Y., 2013. Prognostic significance of AKT/mTOR and MAPK pathways and antitumor effect of mTOR inhibitor in NF1-related and sporadic malignant peripheral nerve sheath tumors. Clin Cancer Res 19, 450–461. 10.1158/1078-0432.CCR-12-1067

Evans, D.G., Howard, E., Giblin, C., Clancy, T., Spencer, H., Huson, S.M., Lalloo, F., 2010. Birth incidence and prevalence of tumor-prone syndromes: estimates from a UK family genetic register service. Am J Med Genet A 152A, 327–332. 10.1002/ajmg.a.33139

Fletcher, P.J., Selhi, Z.F., Azampanah, A., Sills, T.L., 2001. Reduced brain serotonin activity disrupts prepulse inhibition of the acoustic startle reflex. Effects of 5,7-dihydroxytryptamine and p-chlorophenylalanine. Neuropsychopharmacology 24, 399–409. 10.1016/S0893-133X(00)00215-3

Förster, D., Arnold-Ammer, I., Laurell, E., Barker, A.J., Fernandes, A.M., Finger-Baier, K., Filosa, A., Helmbrecht, T.O., Kölsch, Y., Kühn, E., Robles, E., Slanchev, K., Thiele, T.R., Baier, H., Kubo, F., 2017. Genetic targeting and anatomical registration of neuronal populations in the zebrafish brain with a new set of BAC transgenic tools. Sci Rep 7, 5230. 10.1038/s41598-017-04657-x

Giovannitti, J.A., Thoms, S.M., Crawford, J.J., 2015. Alpha-2 adrenergic receptor agonists: a review of current clinical applications. Anesth Prog 62, 31–39. 10.2344/0003-3006-62.1.31

Guo, H.F., The, I., Hannan, F., Bernards, A., Zhong, Y., 1997. Requirement of Drosophila NF1 for activation of adenylyl cyclase by PACAP38-like neuropeptides. Science 276, 795–798. 10.1126/science.276.5313.795

Hannan, F., Ho, I., Tong, J.J., Zhu, Y., Nurnberg, P., Zhong, Y., 2006. Effect of neurofibromatosis type I mutations on a novel pathway for adenylyl cyclase activation requiring neurofibromin and Ras. Hum Mol Genet 15, 1087–1098. 10.1093/hmg/ddl023

Hecker, A., Schulze, W., Oster, J., Richter, D.O., Schuster, S., 2020. Removing a single neuron in a vertebrate brain forever abolishes an essential behavior. Proc Natl Acad Sci U S A 117, 3254– 3260. 10.1073/pnas.1918578117

Ho, I.S., Hannan, F., Guo, H.-F., Hakker, I., Zhong, Y., 2007. Distinct functional domains of neurofibromatosis type 1 regulate immediate versus long-term memory formation. J Neurosci 27, 6852–6857. 10.1523/JNEUROSCI.0933-07.2007

Huang, H.-S., Allen, J.A., Mabb, A.M., King, I.F., Miriyala, J., Taylor-Blake, B., Sciaky, N., Dutton, J.W., Lee, H.-M., Chen, X., Jin, J., Bridges, A.S., Zylka, M.J., Roth, B.L., Philpot, B.D., 2011. Topoisomerase inhibitors unsilence the dormant allele of Ube3a in neurons. Nature 481, 185–189. 10.1038/nature10726

Huang, H.-S., Burns, A.J., Nonneman, R.J., Baker, L.K., Riddick, N.V., Nikolova, V.D., Riday, T.T., Yashiro, K., Philpot, B.D., Moy, S.S., 2013. Behavioral deficits in an Angelman syndrome model: effects of genetic background and age. Behav Brain Res 243, 79–90. 10.1016/j.bbr.2012.12.052

Huson, S.M., Compston, D.A., Harper, P.S., 1989. A genetic study of von Recklinghausen neurofibromatosis in south east Wales. II. Guidelines for genetic counselling. J Med Genet 26, 712–721. 10.1136/jmg.26.11.712

Hyman, S.L., Shores, A., North, K.N., 2005. The nature and frequency of cognitive deficits in children with neurofibromatosis type 1. Neurology 65, 1037–1044. 10.1212/01.wnl.0000179303.72345.ce

Jain, R.A., Wolman, M.A., Marsden, K.C., Nelson, J.C., Shoenhard, H., Echeverry, F.A., Szi, C., Bell, H., Skinner, J., Cobbs, E.N., Sawada, K., Zamora, A.D., Pereda, A.E., Granato, M., 2018. A Forward Genetic Screen in Zebrafish Identifies the G-Protein-Coupled Receptor CaSR as a Modulator of Sensorimotor Decision Making. Curr Biol 28, 1357–1369.e5. 10.1016/j.cub.2018.03.025

Jiang, H., Lei, R., Ding, S.-W., Zhu, S., 2014. Skewer: a fast and accurate adapter trimmer for next-generation sequencing paired-end reads. BMC Bioinformatics 15, 182. 10.1186/1471-2105-15-182

Johannessen, C.M., Reczek, E.E., James, M.F., Brems, H., Legius, E., Cichowski, K., 2005. The NF1 tumor suppressor critically regulates TSC2 and mTOR. Proc Natl Acad Sci U S A 102, 8573–8578. 10.1073/pnas.0503224102

Karaosmanoglu, B., Kocaefe, Ç.Y., Söylemezoğlu, F., Anlar, B., Varan, A., Vargel, İ., Ayter, S., 2018. Heightened CXCR4 and CXCL12 expression in NF1-associated neurofibromas. Childs Nerv Syst 34, 877–882. 10.1007/s00381-018-3745-6

Kimmel, C.B., Patterson, J., Kimmel, R.O., 1974. The development and behavioral characteristics of the startle response in the zebra fish. Dev Psychobiol 7, 47–60. 10.1002/dev.420070109

Krab, L.C., de Goede-Bolder, A., Aarsen, F.K., Pluijm, S.M.F., Bouman, M.J., van der Geest, J.N., Lequin, M., Catsman, C.E., Arts, W.F.M., Kushner, S.A., Silva, A.J., de Zeeuw, C.I., Moll, H.A., Elgersma, Y., 2008. Effect of simvastatin on cognitive functioning in children with neurofibromatosis type 1: a randomized controlled trial. JAMA 300, 287–294. 10.1001/jama.300.3.287

Krab, L.C., Oostenbrink, R., de Goede-Bolder, A., Aarsen, F.K., Elgersma, Y., Moll, H.A., 2009. Health-related quality of life in children with neurofibromatosis type 1: contribution of demographic factors, disease-related factors, and behavior. J Pediatr 154, 420–425, 425.e1. 10.1016/j.jpeds.2008.08.045

Lavoie, H., Gagnon, J., Therrien, M., 2020. ERK signalling: a master regulator of cell behaviour, life and fate. Nat Rev Mol Cell Biol 21, 607–632. 10.1038/s41580-020-0255-7

Li, B., Dewey, C.N., 2011. RSEM: accurate transcript quantification from RNA-Seq data with or without a reference genome. BMC Bioinformatics 12, 323. 10.1186/1471-2105-12-323

Li, W., Cui, Y., Kushner, S.A., Brown, R.A.M., Jentsch, J.D., Frankland, P.W., Cannon, T.D., Silva, A.J., 2005. The HMG-CoA reductase inhibitor lovastatin reverses the learning and attention deficits in a mouse model of neurofibromatosis type 1. Curr Biol 15, 1961–1967. 10.1016/j.cub.2005.09.043

Liu, K.S., Fetcho, J.R., 1999. Laser ablations reveal functional relationships of segmental hindbrain neurons in zebrafish. Neuron 23, 325–335. 10.1016/s0896-6273(00)80783-7

Mainberger, F., Jung, N.H., Zenker, M., Wahlländer, U., Freudenberg, L., Langer, S., Berweck, S., Winkler, T., Straube, A., Heinen, F., Granström, S., Mautner, V.-F., Lidzba, K., Mall, V., 2013. Lovastatin improves impaired synaptic plasticity and phasic alertness in patients with neurofibromatosis type 1. BMC Neurol 13, 131. 10.1186/1471-2377-13-131

Marchuk, D.A., Saulino, A.M., Tavakkol, R., Swaroop, M., Wallace, M.R., Andersen, L.B., Mitchell, A.L., Gutmann, D.H., Boguski, M., Collins, F.S., 1991. cDNA cloning of the type 1 neurofibromatosis gene: complete sequence of the NF1 gene product. Genomics 11, 931–940. 10.1016/0888-7543(91)90017-9

Marsden, K.C., Jain, R.A., Wolman, M.A., Echeverry, F.A., Nelson, J.C., Hayer, K.E., Miltenberg, B., Pereda, A.E., Granato, M., 2018. A Cyfip2-Dependent Excitatory Interneuron Pathway Establishes the Innate Startle Threshold. Cell Rep 23, 878–887. 10.1016/j.celrep.2018.03.095

Miller, A.H., Halloran, M.C., 2022. Mechanistic insights from animal models of neurofibromatosis type 1 cognitive impairment. Dis Model Mech 15, dmm049422. 10.1242/dmm.049422

Mo, W., Chen, J., Patel, A., Zhang, L., Chau, V., Li, Y., Cho, W., Lim, K., Xu, J., Lazar, A.J., Creighton, C.J., Bolshakov, S., McKay, R.M., Lev, D., Le, L.Q., Parada, L.F., 2013. CXCR4/CXCL12 mediate autocrine cell-cycle progression in NF1-associated malignant peripheral nerve sheath tumors. Cell 152, 1077–1090. 10.1016/j.cell.2013.01.053

Padmanabhan, A., Lee, J.-S., Ismat, F.A., Lu, M.M., Lawson, N.D., Kanki, J.P., Look, A.T., Epstein, J.A., 2009. Cardiac and vascular functions of the zebrafish orthologues of the type I neurofibromatosis gene NFI. Proc Natl Acad Sci U S A 106, 22305–22310. 10.1073/pnas.0901932106

Payne, J.M., Barton, B., Ullrich, N.J., Cantor, A., Hearps, S.J.C., Cutter, G., Rosser, T., Walsh, K.S., Gioia, G.A., Wolters, P.L., Tonsgard, J., Schorry, E., Viskochil, D., Klesse, L., Fisher, M., Gutmann, D.H., Silva, A.J., Hunter, S.J., Rey-Casserly, C., Cantor, N.L., Byars, A.W., Stavinoha, P.L., Ackerson, J.D., Armstrong, C.L., Isenberg, J., O’Neil, S.H., Packer, R.J., Korf, B., Acosta, M.T., North, K.N., NF Clinical Trials Consortium, 2016. Randomized placebo-controlled study of lovastatin in children with neurofibromatosis type 1. Neurology 87, 2575–2584. 10.1212/WNL.0000000000003435

Randlett, O., Haesemeyer, M., Forkin, G., Shoenhard, H., Schier, A.F., Engert, F., Granato, M., 2019. Distributed Plasticity Drives Visual Habituation Learning in Larval Zebrafish. Curr Biol 29, 1337–1345.e4. 10.1016/j.cub.2019.02.039

Randlett, O., Wee, C.L., Naumann, E.A., Nnaemeka, O., Schoppik, D., Fitzgerald, J.E., Portugues, R., Lacoste, A.M.B., Riegler, C., Engert, F., Schier, A.F., 2015. Whole-brain activity mapping onto a zebrafish brain atlas. Nat Methods 12, 1039–1046. 10.1038/nmeth.3581

Reiner, A., Yekutieli, D., Benjamini, Y., 2003. Identifying differentially expressed genes usingfalse discovery rate controlling procedures. Bioinformatics 19, 368–375. 10.1093/bioinformatics/btf877

Rigdon, G.C., Weatherspoon, J.K., 1992. 5-Hydroxytryptamine 1a receptor agonists block prepulse inhibition of acoustic startle reflex. J Pharmacol Exp Ther 263, 486–493.

Robinson, M.D., McCarthy, D.J., Smyth, G.K., 2010. edgeR : a Bioconductor package for differential expression analysis of digital gene expression data. Bioinformatics 26, 139–140. 10.1093/bioinformatics/btp616

Robinson, M.D., Oshlack, A., 2010. A scaling normalization method for differential expression analysis of RNA-seq data. Genome Biol 11, R25. 10.1186/gb-2010-11-3-r25

Rosen, L.B., Ginty, D.D., Weber, M.J., Greenberg, M.E., 1994. Membrane depolarization and calcium influx stimulate MEK and MAP kinase via activation of Ras. Neuron 12, 1207–1221. 10.1016/0896-6273(94)90438-3

Ruuskanen, J.O., Peitsaro, N., Kaslin, J.V.M., Panula, P., Scheinin, M., 2005. Expression and function of alpha-adrenoceptors in zebrafish: drug effects, mRNA and receptor distributions. J Neurochem 94, 1559–1569. 10.1111/j.1471-4159.2005.03305.x

Schindelin, J., Arganda-Carreras, I., Frise, E., Kaynig, V., Longair, M., Pietzsch, T., Preibisch, S., Rueden, C., Saalfeld, S., Schmid, B., Tinevez, J.-Y., White, D.J., Hartenstein, V., Eliceiri, K., Tomancak, P., Cardona, A., 2012. Fiji: an open-source platform for biological-image analysis. Nat Methods 9, 676–682. 10.1038/nmeth.2019

Schmitt, J.M., Stork, P.J.S., 2002. PKA Phosphorylation of Src Mediates cAMP’s Inhibition of Cell Growth via Rap1. Molecular Cell 9, 85–94. 10.1016/S1097-2765(01)00432-4

Shilyansky, C., Karlsgodt, K.H., Cummings, D.M., Sidiropoulou, K., Hardt, M., James, A.S., Ehninger, D., Bearden, C.E., Poirazi, P., Jentsch, J.D., Cannon, T.D., Levine, M.S., Silva, A.J., 2010. Neurofibromin regulates corticostriatal inhibitory networks during working memory performance. Proc Natl Acad Sci U S A 107, 13141–13146. 10.1073/pnas.1004829107

Shin, J., Padmanabhan, A., de Groh, E.D., Lee, J.-S., Haidar, S., Dahlberg, S., Guo, F., He, S., Wolman, M.A., Granato, M., Lawson, N.D., Wolfe, S.A., Kim, S.-H., Solnica-Krezel, L., Kanki, J.P., Ligon, K.L., Epstein, J.A., Look, A.T., 2012. Zebrafish neurofibromatosis type 1 genes have redundant functions in tumorigenesis and embryonic development. Dis Model Mech 5, 881–894. 10.1242/dmm.009779

Sipes, T.A., Geyer, M.A., 1994. Multiple serotonin receptor subtypes modulate prepulse inhibition of the startle response in rats. Neuropharmacology 33, 441–448. 10.1016/0028-3908(94)90074-4

Stefan, E., Malleshaiah, M.K., Breton, B., Ear, P.H., Bachmann, V., Beyermann, M., Bouvier, M., Michnick, S.W., 2011. PKA regulatory subunits mediate synergy among conserved G-protein-coupled receptor cascades. Nat Commun 2, 598. 10.1038/ncomms1605

Stivaros, S., Garg, S., Tziraki, M., Cai, Y., Thomas, O., Mellor, J., Morris, A.A., Jim, C., Szumanska-Ryt, K., Parkes, L.M., Haroon, H.A., Montaldi, D., Webb, N., Keane, J., Castellanos, F.X., Silva, A.J., Huson, S., Williams, S., Gareth Evans, D., Emsley, R., Green, J., SANTA Consortium, 2018. Randomised controlled trial of simvastatin treatment for autism in young children with neurofibromatosis type 1 (SANTA). Mol Autism 9, 12. 10.1186/s13229-018-0190-z

Stumm, R.K., Rummel, J., Junker, V., Culmsee, C., Pfeiffer, M., Krieglstein, J., Höllt, V., Schulz, S., 2002. A dual role for the SDF-1/CXCR4 chemokine receptor system in adult brain: isoform-selective regulation of SDF-1 expression modulates CXCR4-dependent neuronal plasticity and cerebral leukocyte recruitment after focal ischemia. J Neurosci 22, 5865–5878. 10.1523/JNEUROSCI.22-14-05865.2002

Subhash, S., Kanduri, C., 2016. GeneSCF: a real-time based functional enrichment tool with support for multiple organisms. BMC Bioinformatics 17, 365. 10.1186/s12859-016-1250-z

Taskén, K., Aandahl, E.M., 2004. Localized effects of cAMP mediated by distinct routes of protein kinase A. Physiol Rev 84, 137–167. 10.1152/physrev.00021.2003

Thomas, G.M., Huganir, R.L., 2004. MAPK cascade signalling and synaptic plasticity. Nat Rev Neurosci 5, 173–183. 10.1038/nrn1346

Tong, J., Hannan, F., Zhu, Y., Bernards, A., Zhong, Y., 2002. Neurofibromin regulates G protein-stimulated adenylyl cyclase activity. Nat Neurosci 5, 95–96. 10.1038/nn792

Torres Nupan, M.M., Velez Van Meerbeke, A., López Cabra, C.A., Herrera Gomez, P.M., 2017. Cognitive and Behavioral Disorders in Children with Neurofibromatosis Type 1. Front Pediatr 5, 227. 10.3389/fped.2017.00227

Uusitalo, E., Leppävirta, J., Koffert, A., Suominen, S., Vahtera, J., Vahlberg, T., Pöyhönen, M., Peltonen, J., Peltonen, S., 2015. Incidence and mortality of neurofibromatosis: a total population study in Finland. J Invest Dermatol 135, 904–906. 10.1038/jid.2014.465

van der Vaart, T., Plasschaert, E., Rietman, A.B., Renard, M., Oostenbrink, R., Vogels, A., de Wit, M.-C.Y., Descheemaeker, M.-J., Vergouwe, Y., Catsman-Berrevoets, C.E., Legius, E., Elgersma, Y., Moll, H.A., 2013. Simvastatin for cognitive deficits and behavioural problems in patients with neurofibromatosis type 1 (NF1-SIMCODA): a randomised, placebo-controlled trial. Lancet Neurol 12, 1076–1083. 10.1016/S1474-4422(13)70227-8

Vanwalleghem, G., Heap, L.A., Scott, E.K., 2017. A profile of auditory-responsive neurons in the larval zebrafish brain. J Comp Neurol 525, 3031–3043. 10.1002/cne.24258

Varni, J.W., Nutakki, K., Swigonski, N.L., 2019. Pain, skin sensations symptoms, and cognitive functioning predictors of health-related quality of life in pediatric patients with Neurofibromatosis Type 1. Qual Life Res 28, 1047–1052. 10.1007/s11136-018-2055-5

Walker, J.A., Tchoudakova, A.V., McKenney, P.T., Brill, S., Wu, D., Cowley, G.S., Hariharan, I.K., Bernards, A., 2006. Reduced growth of Drosophila neurofibromatosis 1 mutants reflects a non-cell-autonomous requirement for GTPase-Activating Protein activity in larval neurons. Genes Dev 20, 3311–3323. 10.1101/gad.1466806

Walsh, K.S., Wolters, P.L., Widemann, B.C., Del Castillo, A., Sady, M.D., Inker, T., Roderick, M.C., Martin, S., Toledo-Tamula, M.A., Struemph, K., Paltin, I., Collier, V., Mullin, K., Fisher, M.J., Packer, R.J., 2021. Impact of MEK Inhibitor Therapy on Neurocognitive Functioning in NF1. Neurol Genet 7, e616. 10.1212/NXG.0000000000000616

Warrington, N.M., Woerner, B.M., Daginakatte, G.C., Dasgupta, B., Perry, A., Gutmann, D.H., Rubin, J.B., 2007. Spatiotemporal differences in CXCL12 expression and cyclic AMP underlie the unique pattern of optic glioma growth in neurofibromatosis type 1. Cancer Res 67, 8588–8595. 10.1158/0008-5472.CAN-06-2220

Wolman, M.A., de Groh, E.D., McBride, S.M., Jongens, T.A., Granato, M., Epstein, J.A., 2014. Modulation of cAMP and ras signaling pathways improves distinct behavioral deficits in a zebrafish model of neurofibromatosis type 1. Cell Rep 8, 1265–1270. 10.1016/j.celrep.2014.07.054

Wolman, M.A., Jain, R.A., Liss, L., Granato, M., 2011. Chemical modulation of memory formation in larval zebrafish. Proc Natl Acad Sci U S A 108, 15468–15473. 10.1073/pnas.1107156108

Wolman, M.A., Jain, R.A., Marsden, K.C., Bell, H., Skinner, J., Hayer, K.E., Hogenesch, J.B., Granato, M., 2015. A genome-wide screen identifies PAPP-AA-mediated IGFR signaling as a novel regulator of habituation learning. Neuron 85, 1200–1211. 10.1016/j.neuron.2015.02.025

