## Supplementary data for "Inhibition of Cxcr4 chemokine receptor signaling improves habituation learning and increases cAMP-PKA signaling in a zebrafish model of Neurofibromatosis type 1"

**Supplementary Figures and Tables**

**Supplementary Figure 1.**

**(A-B)** Histograms demonstrating the frequency distribution of (A) SLC responses and (B) total reactions to both low and high-intensity stimuli averaged across experimental days in 5 dpf non-treated *nf1* mutants. Each average was recorded from groups of 32 *nf1* mutant larvae. A nonlinear regression was fit to each distribution. The distribution average and standard deviation was used to calculate Z-scores to determine compounds that significantly affect behavioral measures. **(C)** Histograms demonstrating the frequency distribution of Z-scores calculated for effects of individual compounds on total reactions to (Top) low-intensity stimuli and (Bottom) high-intensity stimuli. A Z-score threshold (2.326) was set representing the one-sided 99% confidence interval. The number of compounds identified are indicated on the graphs. Each compound was tested on a group of 32 *nf1* mutant larvae. **(D)** Habituation ± SEM for 5 dpf wild-type and *nf1* mutant larvae treated with Plerixafor 8 HCl. Data points represent average habituation from all larvae tested within each genotype at each treatment dose (sample sizes: WT control n=29, 300nM n=26, 1µM n=27, 3µM n=18; $nf1a^{+/-};nf1b^{+/-}$ control n=22, 300nM n=14, 1µM n=15, 3µM n=9; $nf1a^{+/-};nf1b^{-/-}$ control n=17, 300nM n=14, 1µM n=14, 3µM n=18; $nf1a^{-/-};nf1b^{+/-}$ control n=16, 300nM n=16, 1µM n=16, 3µM n=9; $nf1a^{-/-};nf1b^{-/-}$ control n=22, 300nM n=24, 1µM n=16, 3µM n=9). Two-way ANOVA showed statistically significant differences between treatment groups ($F(3,331) = 13.90$, $p<0.001$), as well as a statistically significant interaction between genotype and treatment factors ($F(12,331) = 2.260$, $p = 0.009$). Dunnet's adjusted p-values (below graphs) were used for comparing non-treated larvae within each genotype and comparisons to wild-type larvae within each treatment dose. **(E)** Percent initiation of total reactions ± SEM in 5 dpf wild-type and *nf1* mutant larvae treated with Plerixafor 8HCl. Data points represent average initiation percentage from all larvae tested within each genotype at each treatment dose (sample sizes: WT control n=32, 300nM n=32, 1µM n=32, 3µM n=30; $nf1a^{+/-};nf1b^{+/-}$ control n=24, 300nM n=17, 1µM n=21, 3µM n=17; $nf1a^{+/-};nf1b^{-/-}$ control n=17, 300nM n=17, 1µM n=18, 3µM n=26; $nf1a^{-/-};nf1b^{+/-}$ control n=17, 300nM n=17, 1µM n=21, 3µM n=17; $nf1a^{-/-};nf1b^{-/-}$ control n=22, 300nM n=26, 1µM n=20, 3µM n=11). Two-way ANOVA showed statistically significant differences between treatment groups ($F(3,414) = 11.27$, $p<0.001$. Dunnet's adjusted p-values (below graphs) were used for comparisons between non-treated larvae within each genotype

### Supp. Fig 1

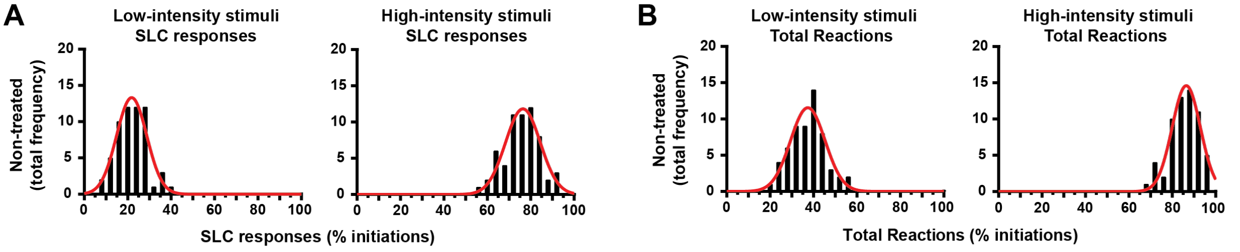


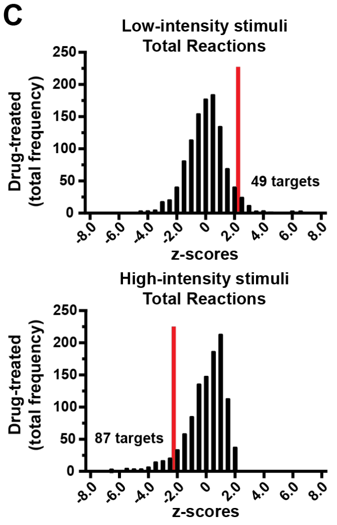


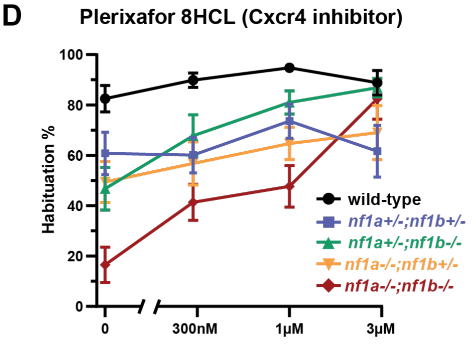


|  | 300nM | 1μM | 3μM |
|---|---|---|---|
| wild-type | ns | ns | ns |
| *nf1a+/- nf1b+/-* | ns | ns | ns |
| *nf1a+/- nf1b-/-* | ns | <0.01 | <0.01 |
| *nf1a-/- nf1b+/-* | ns | ns | ns |
| *nf1a-/- nf1b-/-* | <0.01 | <0.01 | <0.01 |

Comparisons to non-treated larvae

|  | 0nM | 300nM | 1μM | 3μM |
|---|---|---|---|---|
| *nf1a+/- nf1b+/-* | 0.02 | <0.01 | ns | ns |
| *nf1a+/- nf1b-/-* | <0.01 | ns | ns | ns |
| *nf1a-/- nf1b+/-* | <0.01 | <0.01 | <0.01 | ns |
| *nf1a-/- nf1b-/-* | <0.01 | <0.01 | <0.01 | ns |

Comparisons to wild-type larvae

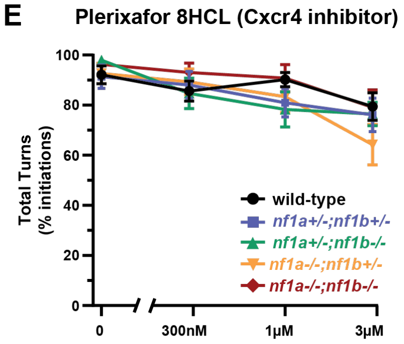


|  | 300nM | 1μM | 3μM |
|---|---|---|---|
| wild-type | ns | ns | ns |
| *nf1a+/- nf1b+/-* | ns | ns | ns |
| *nf1a+/- nf1b-/-* | ns | 0.03 | <0.01 |
| *nf1a-/- nf1b+/-* | ns | ns | <0.01 |
| *nf1a-/- nf1b-/-* | ns | ns | ns |

Comparisons to non-treated larvae

**Supplementary Table 1. Individual compounds that regulate acoustically evoked behaviors in *nf1* mutant larvae**

List of small-molecule compounds identified as targets in Figure 1. Compound names and Biological Targets were defined by the manufacturer (Selleck). Z-scores that surpassed the one-sided 99% confidence interval are highlighted in the respective columns.

| Compound | Biological Target | Low-intensity Total Reaction (z-score) | High-intensity Total Reaction (z-score) | Habituation stimuli 41-50 (z-score) | Habituation stimuli 51-60 (z-score) | Prepulse Inhibition (z-score) |
|---|---|---|---|---|---|---|
| Acebutolol HCl | Adrenergic Receptor | 0.421308 | 0.281543 | 1.915521 | 1.366191 | 2.558511 |
| Acitretin | Retinoid Receptor | 3.978182 | -2.36597 | -11.4933 | -7.72817 | -17.8239 |
| Aclidinium Bromide | AChR | -0.94632 | -0.69273 | 0.988253 | 2.835919 | 1.523516 |
| Allopurinol (Zyloprim) | ROS | -1.68461 | -1.30177 | 1.761356 | 2.651828 | 0.72008 |
| Alprostadil (Caverject) | Immunology & Inflammation related | -4.29393 | -10.1228 | 2.495002 | 2.176356 | 3.696938 |
| Alverine Citrate | OX Receptor | -2.03173 | -3.70241 | -0.38296 | -0.15575 | 0.954885 |
| Amantadine hydrochloride (Symmetrel) | Dopamine Receptor | 2.826292 | 1.227472 | -1.27919 | -0.25266 | 0.271315 |
| Amfenac Sodium (monohydrate) | COX | 2.537266 | 1.302105 | -0.34468 | 0.687782 | -0.35063 |
| AMG-073 HCl (Cinacalcet hydrochloride) | Others | 6.214039 | 2.188712 | -3.66821 | -2.3552 | -4.20904 |
| Amiloride hydrochloride (Midamor) | Others | 2.478555 | 1.265751 | -0.70448 | -0.02291 | -0.2993 |
| Amitriptyline HCl | 5-HT Receptor | -2.74438 | -7.09409 | 2.631288 | 0.491604 | 3.20173 |
| Amorolfine Hydrochloride | Anti-infection | -2.30251 | -4.59723 | 2.367701 | 2.341202 | 0.605162 |
| Amoxapine | GlyT | 0.632751 | 1.260477 | 2.543958 | 0.981392 | 0.847341 |
| Ampiroxicam | COX | 0.603552 | 0.095663 | 0.807294 | 1.152838 | 2.613325 |
| Arecoline | AChR | 3.01266 | 2.015518 | 0.333387 | 0.437928 | 1.057253 |
| Aripiprazole (Abilify) | 5-HT Receptor | 0.020134 | 0.282315 | 1.338088 | 2.433811 | 0.992232 |
| Asenapine | Others | 3.021246 | 1.88583 | -2.45118 | -0.95165 | -1.86042 |
| Atomoxetine HCl | 5-HT Receptor | -3.99261 | -6.39459 | 2.160169 | 2.605673 | 3.294274 |
| Atorvastatin calcium (Lipitor) | HMG-CoA Reductase | -0.09052 | -0.89511 | 0.761225 | 1.197723 | 2.47769 |
| Atracurium besylate | AChR | -2.41247 | -4.53948 | -1.89733 | 1.262034 | 0.486342 |
| Azacitidine (Vidaza) | DNA Methyltransferase | -0.64421 | -3.26243 | 0.062856 | -0.66749 | -0.27474 |
| Azelnidipine | Calcium Channel | -2.07436 | -2.76495 | 0.722604 | 1.510537 | 1.522512 |

| Compound | Biological Target | Low-intensity Total Reaction (z-score) | High-intensity Total Reaction (z-score) | Habituation stimuli 41-50 (z-score) | Habituation stimuli 51-60 (z-score) | Prepulse Inhibition (z-score) |
|---|---|---|---|---|---|---|
| Azilsartan Medoxomil (TAK-491) | RAAS | -1.93226 | -2.77738 | 0.061098 | 0.577246 | 1.395656 |
| Azlocillin sodium salt | Anti-infection | -2.00674 | -2.70673 | -0.51041 | 0.51255 | 2.255144 |
| Benidipine hydrochloride | Calcium Channel | -2.67413 | -3.95266 | 1.166776 | 1.236766 | 1.358876 |
| Benzbromarone | P450 (e.g. CYP17) | -3.01591 | -5.26609 | 1.792406 | 2.24619 | 0.145903 |
| Benzocaine | Sodium Channel | 2.366598 | 1.35541 | 0.546085 | 0.086273 | 0.767483 |
| Benztropine mesylate | Dopamine Receptor | -0.3416 | 0.213576 | 2.607055 | 1.907421 | 1.072365 |
| Besifloxacin HCl (Besivance) | Anti-infection | -1.87377 | -2.76312 | 2.032134 | 1.928163 | 2.388905 |
| Betamethasone (Celestone) | Glucocorticoid Receptor | -0.07529 | -0.31518 | -0.2989 | 0.499749 | 2.653669 |
| BIBR 953 (Dabigatran etexilate, Pradaxa) | Anti-infection | -1.90987 | -3.18777 | 0.037313 | 0.187195 | 1.568575 |
| Bifonazole | Anti-infection | 0.652738 | -3.90904 | 2.608574 | 2.644551 | -1.16206 |
| Bleomycin sulfate | DNA/RNA Synthesis | -1.3496 | -2.48893 | 0.408625 | 0.694112 | 1.023436 |
| Bumetanide | Others | -1.55914 | -2.40896 | 0.133825 | -0.12731 | -0.43519 |
| Calcitriol (Rocaltrol) | Vitamin | -3.11756 | -5.47654 | 0.812808 | 1.21524 | -0.39294 |
| Carvedilol | DNA/RNA Synthesis | -1.88249 | -2.62676 | -1.81935 | 2.124067 | 1.087218 |
| Caspofungin acetate | DNA/RNA Synthesis | -2.81052 | -3.95944 | -1.82237 | -1.65394 | 1.1341 |
| Celecoxib | COX | -2.94081 | -3.75779 | 2.150513 | 2.191325 | -0.71837 |
| Chloramphenicol (Chloromycetin) | Anti-infection | 2.384046 | 1.295524 | 1.296416 | 1.139134 | 1.692065 |
| Chlorocresol | Calcium Channel | 2.333651 | 1.057426 | -1.65341 | -0.91732 | 0.050184 |
| Chloroxine | Anti-infection | 2.899993 | 1.74467 | 1.205407 | 2.112354 | -2.64596 |
| Chlorpromazine (Sonazine) | Potassium Channel, Dopamine Receptor | -1.82025 | -0.28737 | 1.820779 | 3.118333 | 1.506288 |
| Chlorpropamide | Others | -0.78162 | -0.55512 | 1.556138 | 1.531455 | 2.540194 |
| Chlortetracycline HCl | Anti-infection | -0.00549 | -0.65544 | 1.155105 | 2.328873 | 1.20366 |
| Chlorzoxazone | P450 (e.g. CYP17) | -0.73525 | -0.80768 | 2.320023 | 2.471749 | 1.489087 |
| Cilnidipine | Calcium Channel | -2.63704 | -7.11322 | -2.21013 | 0.576538 | 2.349453 |
| Cleviprex (Clevidipine) | Calcium Channel | 0.0789 | 0.010525 | 1.593188 | 2.395967 | 2.125829 |
| Clomipramine hydrochloride (Anafranil) | 5-HT Receptor | -2.97173 | -3.15071 | 1.435527 | 1.895656 | 0.866145 |
| Closantel | Anti-infection | -3.04217 | -6.23522 | 1.975814 | 3.086274 | 3.014535 |
| Crystal violet | Others | -4.10235 | -6.32222 | -13.2203 | -9.13011 | -0.38262 |

| Compound | Biological Target | Low-intensity Total Reaction (z-score) | High-intensity Total Reaction (z-score) | Habituation stimuli 41-50 (z-score) | Habituation stimuli 51-60 (z-score) | Prepulse Inhibition (z-score) |
|---|---|---|---|---|---|---|
| Deferasirox (Exjade) | P450 (e.g. CYP17) | -3.09852 | -5.01799 | -0.04602 | 0.327068 | 1.466917 |
| Deoxycorticosterone acetate | Adrenergic Receptor | 2.52218 | 1.834077 | 0.544899 | 1.066751 | -1.62806 |
| Detomidine HCl | Adrenergic Receptor | -1.3203 | -2.62053 | 2.455064 | 2.773297 | 1.137442 |
| Dexmedetomidine HCl (Precedex) | Adrenergic Receptor | 0.96478 | 0.130373 | 2.160723 | 2.512874 | 1.758576 |
| Dextrose (D-glucose) | NA | 2.619867 | 1.745691 | 0.483317 | 0.248384 | 0.174683 |
| Dibenzothiophene | NA | 2.618149 | 0.800315 | -0.22881 | -0.17572 | -1.45731 |
| Dichlorisone Acetate | NA | 3.039094 | 2.01688 | -1.71742 | -1.17535 | -1.29533 |
| Dicyclomine HCl | Others | -1.93242 | -2.43647 | -0.32131 | -0.09384 | -0.35077 |
| Diethylstilbestrol (Stilbestrol) | Estrogen / progestogen Receptor | -3.44885 | -9.20853 | 1.28511 | 3.434416 | -0.27641 |
| Difluprednate | Others | 1.073041 | -0.41123 | 2.291753 | 1.96536 | 2.609367 |
| Diperodon HCl | Others | 2.626425 | 1.638508 | -0.40898 | -0.22617 | -0.93076 |
| Doxercalciferol (Hectorol) | Vitamin | -2.77639 | -2.72601 | 1.548088 | 1.544834 | 0.880856 |
| Doxofylline | PDE | -0.44988 | 0.690879 | 1.457308 | 2.468547 | 1.978805 |
| Drospirenone | Estrogen / progestogen Receptor | -1.50385 | -3.58955 | 1.087269 | -0.41493 | 0.093481 |
| Duloxetine HCl (Cymbalta) | 5-HT Receptor | -1.15512 | -3.57433 | 2.273265 | 0.350268 | 1.695556 |
| Dutasteride | 5-alpha Reductase | -2.39773 | -5.01038 | -0.2191 | 0.078269 | 0.901781 |
| Dyclonine HCl | Sodium Channel | -0.16281 | 0.060628 | 2.539832 | 2.09776 | 1.587401 |
| Enoxacin (Penetrex) | Topoisomerase | -2.33652 | -2.87116 | 2.359797 | 2.072229 | 2.487949 |
| Enrofloxacin | Anti-infection | -2.95031 | -3.74461 | 0.816555 | 2.298714 | 1.806858 |
| Epinephrine bitartrate (Adrenalinium) | Adrenergic Receptor | 4.151504 | 1.951379 | 2.081548 | 2.349438 | 0.854977 |
| Erlotinib HCl | EGFR | -2.4701 | -3.72316 | -2.84772 | 0.324386 | 0.514782 |
| Escitalopram oxalate | 5-HT Receptor | -0.20244 | -0.26968 | 0.141512 | 0.106302 | 2.350175 |
| Estradiol | Estrogen / progestogen Receptor | -4.41015 | -10.4013 | 3.376039 | 2.498589 | 3.153063 |
| Ethacridine lactate monohydrate | Anti-infection | -2.47063 | -2.61963 | -0.36537 | 0.279382 | -0.79685 |
| Ethoxzolamide | Others | -1.46991 | -2.59072 | 1.301666 | 0.143869 | 0.344243 |
| Etomidate | GABA Receptor | 6.533383 | 2.283645 | -1.83113 | -1.8623 | -5.77916 |
| Etoposide (VP-16) | Topoisomerase | - 1.361860 | - 2.613320 | 0.407693 | -0.688181 | 1.813234 |
| Evista (Raloxifene Hydrochloride) | mTOR | -1.96727 | -3.25789 | -0.34413 | 0.644483 | 0.681027 |
| Felbamate | NMDAR | 2.423903 | 1.464847 | -3.5872 | -2.71284 | -2.24664 |

| Compound | Biological Target | Low-intensity Total Reaction (z-score) | High-intensity Total Reaction (z-score) | Habituation stimuli 41-50 (z-score) | Habituation stimuli 51-60 (z-score) | Prepulse Inhibition (z-score) |
|---|---|---|---|---|---|---|
| Fenoprofen calcium | Immunology & Inflammation related | 2.419646 | 1.384945 | -2.54282 | -3.20058 | -1.31771 |
| Finasteride | 5-alpha Reductase | -1.45961 | -2.87047 | -1.83036 | -2.24565 | 1.165341 |
| Fluconazole | P450 (e.g. CYP17) | -0.08669 | 1.307592 | 2.046619 | 2.222052 | 2.814241 |
| Flumazenil | GABA Receptor | 2.512103 | 1.696075 | -2.51948 | -1.72657 | -1.69189 |
| Fluvoxamine maleate | 5-HT Receptor | -1.44378 | -0.70475 | 2.335901 | 2.598347 | 0.34965 |
| Ftorafur | NA | -0.69939 | -1.34712 | 0.595654 | 1.070845 | 2.501992 |
| Furosemide (Lasix) | Sodium Channel | 3.073544 | 1.670447 | -1.80722 | -1.19555 | -1.0937 |
| Gabapentin (Neurontin) | GABA Receptor | 0.302814 | -0.28558 | 2.372807 | 1.839923 | 0.569497 |
| Genistein | EGFR, Topoisomerase | -3.37833 | -6.89121 | 2.868168 | 2.956799 | 3.440108 |
| Hyoscyamine (Daturine) | AChR | -0.16349 | -0.65528 | -0.72231 | 0.031917 | 2.328959 |
| Ifosfamide | DNA/RNA Synthesis | -0.59951 | 0.095462 | 1.437829 | 1.889965 | 2.772529 |
| Iloperidone (Fanapt) | 5-HT Receptor | 2.989621 | 2.071364 | -1.7861 | -1.25266 | -0.48867 |
| Imatinib Mesylate | Bcr-Abl, c-Kit, PDGFR | -1.74176 | -0.16099 | 1.517681 | 0.979593 | 2.404743 |
| Imipramine HCl | Others | -2.69516 | -2.53289 | 2.745015 | 2.459705 | 1.554301 |
| Imiquimod | Immunology & Inflammation related | -2.12816 | -3.78363 | -1.37733 | -4.5849 | 1.073567 |
| Irinotecan | Topoisomerase | -1.01973 | -1.98945 | 0.505238 | 0.617193 | 2.425176 |
| Isoxicam | NA | 3.155791 | 2.120319 | -2.59136 | -2.55838 | -2.3253 |
| Isradipine (Dynacirc) | Calcium Channel | -3.92978 | -8.2439 | -1.02488 | 1.500642 | 2.683792 |
| Levamisole Hydrochloride (Ergamisol) | Immunology & Inflammation related | -1.00508 | -2.68452 | -0.69704 | -0.41508 | 1.502713 |
| Levofloxacin (Levaquin) | Topoisomerase | 2.44868 | 0.947148 | -3.01474 | -2.1218 | -1.03826 |
| Levonorgestrel (Levonelle) | Estrogen / progestogen Receptor | -3.44391 | -5.42941 | -1.02342 | -1.03469 | 0.558819 |
| Lomustine (CeeNU) | DNA/RNA Synthesis | -2.68463 | -3.65625 | -3.66974 | -3.46101 | 0.509396 |
| Loratadine | Histamine Receptor | 6.162984 | 2.234647 | -3.42546 | -2.85107 | -5.44635 |
| Malotilate | Others | -2.1258 | -2.44573 | -1.78018 | -2.57236 | -2.0673 |
| MDV3100 (Enzalutamide) | c-Kit, PDGFR | 2.528144 | 1.935217 | -0.94254 | -0.49721 | -1.03652 |
| Medetomidine HCl | Adrenergic Receptor | -1.16295 | -3.05847 | 2.560893 | 2.868119 | 1.855348 |
| Mefenamic acid | COX | 3.583205 | 1.97877 | -2.93854 | -2.86635 | -3.12714 |
| Methscopolamine (Pamine) | AChR | -0.83672 | -0.08322 | 1.735141 | 1.497091 | 2.624781 |
| Mexiletine HCl | Sodium Channel | -1.42073 | -3.03525 | -0.36593 | 0.56033 | 0.828205 |

| Compound | Biological Target | Low-intensity Total Reaction (z-score) | High-intensity Total Reaction (z-score) | Habituation stimuli 41-50 (z-score) | Habituation stimuli 51-60 (z-score) | Prepulse Inhibition (z-score) |
|---|---|---|---|---|---|---|
| Mianserin hydrochloride | Histamine Receptor | 1.098691 | 1.653395 | 2.051582 | 2.698842 | 1.337899 |
| Miconazole nitrate | Anti-infection | -3.05605 | -3.72538 | 2.185008 | 1.694363 | 1.790361 |
| Monobenzone (Benoquin) | Tyrosinase | -1.42947 | -2.94314 | -2.80863 | -1.7141 | 2.662652 |
| Nafamostat mesylate | Serine Protease | -0.33005 | 0.392894 | 1.298217 | 2.586782 | 1.611949 |
| Naftopidil (Flivas) | Adrenergic Receptor | 2.810859 | 1.645313 | 0.385752 | 0.943468 | 0.889912 |
| Netilmicin Sulfate | Anti-infection | -2.59573 | -3.79752 | -0.14516 | 1.683154 | 2.664478 |
| Nilotinib (AMN-107) | Bcr-Abl | -1.88366 | -3.2281 | 0.350868 | 0.857194 | 2.267196 |
| Nimodipine (Nimotop) | Autophagy, Calcium Channel | -4.28226 | -2.81355 | 3.345204 | 2.098739 | -5.50049 |
| Nisoldipine (Sular) | Calcium Channel | -3.64675 | -7.35319 | 3.955216 | 3.561014 | 4.145767 |
| Norfloxacin (Norxacin) | Topoisomerase | -3.17284 | -5.03694 | -0.06236 | 0.536054 | 3.137584 |
| olsalazine sodium | Anti-infection | 0.530279 | 0.769613 | 0.87366 | 0.549903 | 2.343754 |
| Oxybutynin (Ditropan) | AChR | -3.12139 | -2.5005 | 2.652056 | 2.654832 | 0.592397 |
| Oxytetracycline (Terramycin) | Anti-infection | -2.46777 | -2.66433 | 1.645679 | 2.268499 | 1.95702 |
| Paromomycin Sulfate | Anti-infection | -0.90045 | -2.75308 | 0.691035 | 0.593974 | 0.622171 |
| Pazopanib | c-Kit, PDGFR, VEGFR | 2.463096 | 0.986226 | -3.06004 | -2.28175 | -0.84305 |
| Penfluridol | Dopamine Receptor | 0.014331 | 0.023139 | 1.255418 | 2.468923 | 2.183519 |
| Pergolide mesylate | Dopamine Receptor | -2.09512 | -2.79211 | 1.667547 | 1.126088 | 1.533419 |
| Pheniramine Maleate | Histamine Receptor | -3.06354 | -5.48881 | 1.883675 | 1.029556 | 1.240885 |
| Phenylephrine HCl | Adrenergic Receptor | -1.21598 | -0.65 | 1.961588 | 2.741951 | 0.991361 |
| Piromidic Acid | Others | 3.04523 | 1.58124 | -0.78469 | -0.53593 | -0.19587 |
| Piroxicam (Feldene) | COX | -2.8668 | -3.54145 | -3.28321 | -0.87068 | 1.499721 |
| Pitavastatin calcium (Livalo) | HMG-CoA Reductase | -2.12227 | -4.1929 | 1.966497 | 0.600294 | -0.71941 |
| Pizotifen malate | 5-HT Receptor | 2.469186 | 1.761003 | 0.150826 | 0.451528 | 0.392392 |
| Plerixafor (AMD3100) | CXCR | 1.248123 | 0.79636 | 2.337142 | 1.670019 | 2.103862 |
| Pregnenolone | Estrogen / progestogen Receptor | -1.08639 | -2.60407 | 0.204998 | -0.40631 | 0.112481 |
| Procaine (Novocaine) HCl | Anti-infection | -1.08425 | -0.82313 | 0.938866 | 1.39973 | 2.543957 |
| Protionamide (Prothionamide) | Anti-infection | -2.39476 | -3.70332 | -0.69687 | -0.38614 | 0.929215 |
| Pyrithione zinc | Proton Pump, Anti-infection | -3.39147 | -4.738 | -4.10342 | -2.94806 | -0.49392 |

| Compound | Biological Target | Low-intensity Total Reaction (z-score) | High-intensity Total Reaction (z-score) | Habituation stimuli 41-50 (z-score) | Habituation stimuli 51-60 (z-score) | Prepulse Inhibition (z-score) |
|---|---|---|---|---|---|---|
| Ractopamine HCl | Others | 2.972278 | 2.178891 | -0.77249 | -1.30704 | -0.50659 |
| Reboxetine mesylate | Others | -2.37837 | -3.30912 | 1.701515 | 1.775376 | 2.734845 |
| Resveratrol | Autophagy | 2.476726 | 1.983023 | -0.04071 | 0.030355 | -0.64519 |
| Riluzole (Rilutek) | GluR, Sodium Channel | 4.155877 | 1.551304 | 0.454758 | -0.65313 | -5.78648 |
| Rimonabant (SR141716) | Cannabinoid Receptor | 3.757165 | 1.96866 | -4.4425 | -5.22721 | -4.26326 |
| Risedronic acid (Actonel) | NA | -3.10137 | -4.58469 | 3.284599 | 2.297385 | 1.312182 |
| Risperidone (Risperdal) | 5-HT Receptor | -2.09411 | -0.26657 | 2.633452 | 2.715713 | 1.183495 |
| Rivaroxaban (Xarelto) | Factor Xa | 3.728505 | 2.236178 | -0.77368 | -1.45299 | -1.33439 |
| Rizatriptan Benzoate (Maxalt) | 5-HT Receptor | -0.61718 | 0.423092 | 2.194615 | 1.410742 | 2.4724 |
| Roflumilast (Daxas) | PDE | -2.09169 | -3.72878 | -0.54814 | -2.6528 | 3.207306 |
| Rolipram | PDE | -3.11713 | -5.27678 | 1.174834 | 2.54847 | 2.568928 |
| Rolitetracycline | PDE | 2.35482 | 1.104839 | 0.834422 | 0.008613 | 1.536146 |
| Ropivacaine HCl | PDE | -2.03501 | -3.08444 | -0.05032 | 0.931685 | 2.021863 |
| Sarafloxacin HCl | Anti-infection | -1.928 | -3.22107 | -0.53965 | 1.877445 | 1.064599 |
| Sertraline HCl | 5-HT Receptor | -2.81542 | -5.32984 | 1.886099 | 1.873202 | 3.69223 |
| Sitafloxacin hydrate | Anti-infection | -2.3167 | -2.33303 | 1.062152 | 1.513251 | 1.348962 |
| Sodium Gluconate | Others | -0.07384 | 0.004862 | 0.02373 | 0.041292 | 2.41692 |
| Sorafenib (Nexavar) | PDGFR, Raf, VEGFR | -3.22883 | -3.7036 | 1.703912 | 1.91252 | 0.313793 |
| Sotalol (Betapace) | Adrenergic Receptor | 2.358418 | 1.487432 | -2.78704 | -1.96996 | -3.61898 |
| Spectinomycin hydrochloride | Anti-infection | 2.395651 | 1.891663 | -2.45555 | -2.48384 | -2.77414 |
| Sulfamethizole (Proklar) | Anti-infection | -1.63699 | -1.61556 | 1.53491 | 0.567233 | 2.478949 |
| Sumatriptan succinate | 5-HT Receptor | 2.537109 | 1.47144 | -0.91147 | -1.29982 | 0.433707 |
| Sunitinib Malate (Sutent) | c-Kit, PDGFR, VEGFR | -2.27656 | -3.89267 | -1.61475 | -0.34152 | 0.620182 |
| Suprofen (Profenal) | COX | -1.81203 | -2.80394 | 0.945676 | 0.505011 | 0.59218 |
| Tadalafil (Cialis) | PDE | -1.29705 | -1.2088 | 1.378144 | 1.931921 | 3.136495 |
| Tazarotene (Avage) | Retinoid Receptor | 6.296766 | 2.148629 | -4.73147 | -3.96366 | -5.45499 |
| Tebipenem pivoxil (L-084) | Anti-infection | -2.02044 | -2.71412 | -0.76274 | -0.62592 | -0.38115 |
| Terbinafine (Lamisil, Terbinex) | Anti-infection | -2.95611 | -3.66054 | -2.06765 | -1.00166 | 1.720487 |
| Terfenadine | Others | 1.148227 | 0.91325 | -0.91236 | -0.0034 | 2.574038 |
| tetrahydrozoline hydrochloride | Adrenergic Receptor | -2.15227 | -2.89548 | 0.44342 | 1.755001 | 1.079063 |
| Tianeptine sodium | 5-HT Receptor | 3.060531 | 2.08221 | -1.23418 | -1.61037 | -0.88374 |
| tinidazole | Anti-infection | 2.337464 | 1.137173 | -0.6419 | -1.60288 | 0.130478 |
| Tobramycin | Anti-infection | 3.251946 | 1.571647 | 0.591653 | 0.458885 | 0.701805 |

| Compound | Biological Target | Low-intensity Total Reaction (z-score) | High-intensity Total Reaction (z-score) | Habituation stimuli 41-50 (z-score) | Habituation stimuli 51-60 (z-score) | Prepulse Inhibition (z-score) |
|---|---|---|---|---|---|---|
| Tolnaftate | Anti-infection | -3.26407 | -4.96362 | -3.6 | -3.72485 | 0.020652 |
| Toremifene Citrate (Fareston, Acapodene) | Estrogen / progestogen Receptor | 4.742518 | 0.954017 | -3.638 | -2.7311 | -2.45292 |
| Tranexamic acid (Transamin) | Others | -1.62993 | -0.97723 | 2.699371 | 2.458322 | 1.487175 |
| Triclabendazole | Microtubule Associated | -4.2646 | -8.82115 | -0.23189 | -1.56121 | -2.98622 |
| Trilostane | Dehydrogenase | -1.99675 | -3.31741 | -0.73957 | -3.64485 | 0.704744 |
| Valproic acid sodium salt (Sodium valproate) | GABA Receptor, HDAC, Autophagy | -0.67797 | -0.14443 | 1.940246 | 2.090855 | 2.3452 |
| Vardenafil (Vivanza) | PDE | 2.85599 | 1.782779 | -1.61491 | -1.6149 | -1.57769 |
| Varenicline tartrate | AChR | -0.27415 | -0.18556 | 0.985315 | 2.58431 | 1.311162 |
| Vinpocetine (Cavinton) | Sodium Channel | 2.584734 | 0.981705 | -0.2796 | -0.58491 | -0.02913 |
| Vitamin D3 (Cholecalciferol) | NA | 1.052784 | 0.541886 | -0.06757 | 0.071739 | 2.752007 |
| Xylazine HCl | Adrenergic Receptor | 2.582946 | 0.89071 | 0.720803 | 0.350189 | 0.675392 |
| Ziprasidone hydrochloride | 5-HT Receptor, Dopamine Receptor | -0.56489 | 1.050268 | 1.766881 | 2.490617 | 0.833657 |

39
